# Asymmetric Ion Mobility and Interface Displacement Drive the Signal Enhancement in a polymer-electrolyte nanopore

**DOI:** 10.1101/2022.08.17.503612

**Authors:** Fabio Marcuccio, Dimitrios Soulias, Chalmers C. Chau, Sheena E. Radford, Eric W. Hewitt, Paolo Actis, Martin A. Edwards

## Abstract

Solid-state nanopores have been widely employed in the detection of biomolecules, but low signal-to-noise ratios still represent a major obstacle to enable the discrimination of short nucleic acid and protein sequences. The addition of 50% polyethylene glycol (PEG) to the bath solution was recently demonstrated as a simple way to enhance the detection of such biomolecules translocating through a model solid-state nanopore. Here, we provide a comprehensive description of the physics describing a nanopore measurement carried out in 50% PEG that is supported by finite-element modelling and experiments. We demonstrate that the addition of PEG to the external solution introduces a strong imbalance in the transport properties of cations and anions, drastically affecting the characteristic current response of the nanopore. We further show that the strong asymmetric current response is due to a polarity-dependent ion distribution and transport at the nanopipette tip region, leading to either ion depletion or enrichment for few tens of nanometers across the aperture. Under negative potential, when double-stranded DNA molecules translocate, the depleted region (sensing region) significantly improves the sensitivity compared to systems without PEG. We then introduce a displacement of the interface between pore and external solution to simulate the mechanical interactions between analyte and PEG molecules. We found that this displacement affects the ion distribution in the sensing region, enhancing the detection current during the translocation of biomolecules.

## INTRODUCTION

Nanopore sensing is one of the leading label-free techniques for the analysis and manipulation of single molecules due to its high throughput and sensitivity^1–3^. In nanopore measurements, an ionic current is generated by applying a potential between two electrodes situated in two reservoirs separated by a small orifice. In general, the translocation of an analyte through a nanopore causes a decrease in magnitude of the ionic current due to temporary restricted transport of ions across the orifice. However, under low electrolyte concentrations, charged molecules, such as double-stranded DNA (dsDNA), carry a counterion cloud which leads to a local ion enrichment, inducing a current enhancement^1,4,5^. The amplitude, duration and shape of the translocation event provide important information about the physicochemical properties of the molecule, such as size, charge, and shape^1,6^.

Despite the developments in the field over the past decades^7^, using solid-state nanopores to detect proteins and short nucleic acids still remains challenging, requiring nanopores of comparable size to the molecules (< 10 nm diameter), which are difficult to fabricate reproducibly^8^. Furthermore, the nanopore system need to have higher signal-to-noise ratio^9^ to detect small perturbations to the ion current caused by the translocation of molecules, and high bandwidth electronics to characterize rapid translocations with sufficient temporal resolution^10^. Finite element modelling has been extensively used to examine electrokinetic phenomena in nanopores^5,11–14^. In such systems, the ion current is due to the transport of ionic species under the influence of an electric field and its physics can be considerably more complex than that in simple ohmic conductors^15^. For example, the charge on the nanopore wall induces an electric double layer leading to non-uniform ion concentration distributions and the interacting physics of ion transport, electric fields and fluid flows result in a wide range of non-linear behaviour ^12,16,17^.

We have recently reported the enhanced single molecule detection of a nanopore when 50% polyethylene glycol (PEG) is added to the bath solution leading to a 6X increase of the amplitude of the translocation signal^18^.

Here, we describe a mechanism explaining this enhancement by using a combination of experiments and multi-physics modelling^18^. We developed a finite element model by coupling Nernst-Planck, Poisson and Navier-Stokes equations to describe the physics of ion transport under an applied electric field when a nanopore sensing experiment is carried out in presence of 50% PEG. Based on the cation-binding properties of PEG that have been largely discussed in the literature, our model assumes an imbalance between the diffusion coefficients of cations and anions in the bath solution^19–23^. The model reproduces the experimental current-voltage responses in the presence and absence of PEG and provides insight into the ion concentrations and transport rates responsible for the observed behavior. We then provide evidence that a combination of the asymmetric ion mobility at the nanopore and the mechanical interaction between a translocating molecule and the nanopore-bath interface is responsible for the increase in the translocation signals. This new mechanism may inform further developments in nanopore sensing by suggesting that chemical approaches that affect ion mobilities could be used to enhance the sensitivity of the system.

## RESULTS AND DISCUSSION

Figure 1a shows the experimental setup used throughout this work in which a model solid-state nanopore based on a quartz nanopipette (25 nm in diameter) filled with a 0.1 M KCl solution is immersed into a bath containing 0.1 M KCl with or without 50% (w/v) PEG. In nanopore measurements, the current-voltage response characterizes the ion transport, indirectly providing information about the physical properties of the nanopore (size, shape, surface charge). The grey line in Figure 1b shows the current-voltage response of a nanopipette filled with a 0.1M KCl solution and immersed in a bath containing 0.1 M KCl (no PEG). The slightly higher conductivity observed at a positive bias applied vs a negative bias is termed ion-current rectification (ICR) and arises from the negative charge on the glass wall of the nanopipette, which makes the aperture region permselective to cations and this effect has been extensively described in the literature^17,24–26^.

**Figure 1.**
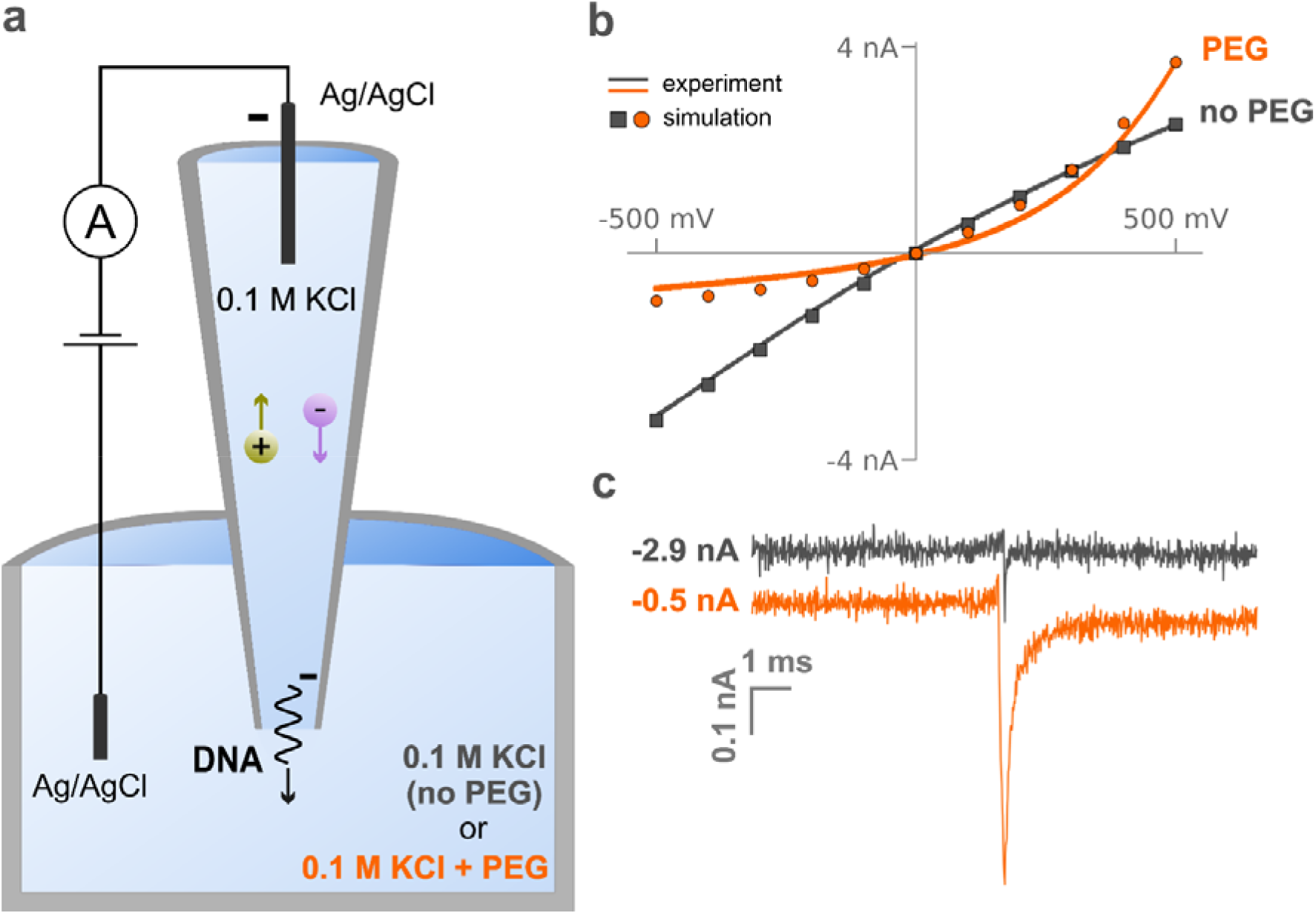
Schematic and representative data for conductive-pulse measurements of double-stranded DNA translocation through a nanopipette. (a) A nanopipette (12.5 nm pore radius), filled with a 10 nM solution of 4.8kbp dsDNA in 0.1 M KCl, is immersed in a solution of the same electrolyte with and without the presence of 50% (w/v) PEG 35K. The application of a negative potential to an Ag/AgCl quasi-reference electrode inside the nanopipette with respect to a ground electrode in the bath solution causes outbound migration of DNA molecules, initially present in the nanopipette solution. (b) Experimental (curves) and simulated (points) voltammograms of the nanopipette in the presence (orange) and absence (black) of PEG in the outside solution. Current trace recorded upon translocation of a dsDNA molecule through the nanopipette aperture with (orange trace) and without (black trace) the presence of PEG in the bath solution.

When the same nanopipette is immersed in a bath of 0.1 M KCl with 50% PEG, a dramatic reversal in the rectification is observed in the *i-V* curve (orange line). The PEG solution is ~9 times less conductive than 0.1 M KCl (Table S1.2, SI1) and, counterintuitively, the ion current observed at +500 mV is greater than the one measured in a PEG-free bath. Also, under negative bias, the ion current is ~4 times lower than observed without PEG in the bath solution. This response cannot be explained only considering the difference in conductivity between the two solutions, or as a rectification effect induced by surface-charge on the nanopore wall, indicating that a fundamentally different nanopore physics is responsible for the observed *i-V* response. We have investigated if the difference in viscosity between the two solution is responsible for the observed experimental difference, but the *i-V* response observed in PEG cannot be reproduced other viscous solutions such as 50% glycerol (S1.6, Supporting Information 1), indicating that viscosity alone cannot be the responsible for such behavior. In the following section, we describe a numerical model of ion transport, from which the calculated ion current (points in Figure 1b) is derived, that explains the anomalous current-voltage response.

As we have previously reported^18^, the presence PEG in the bath solution leads to a 4 fold enhancement of the ion current observed when a single molecule translocates through the nanopore (Figure 1c).

It is worth noting that as the two current traces displayed in Figure 1c were both recorded using the same nanopipette (*r* = 12.5 nm), applied voltage (−500 mV), and composition of the inner solution (0.1 M KCl and 0.1 ng/μl 4.8 kbp dsDNA) and the observed enhancement is only driven by the presence of PEG in the bath solution. In a conventional nanopore measurements where the solution is identical in both reservoirs, the current increase is attributed to the presence of the counter-ion cloud carried by the dsDNA molecule, which results in a temporary increase in the ion concentration in this region, and this effect has been extensively described in the literature^4,5,27,28^. The next section describes a new nanopore physics that not only explains the anomalous *i-V* response, but also the enhanced single molecule sensitivity.

### Finite Element Simulations

We developed a finite element model that coupled ion transport (diffusion, electromigration) and electric fields at different applied potentials. A detailed description of the model is given in the Supporting Information 1 (Section S1) and 2 (COMSOL report). Briefly, a 2D axisymmetric model simulates the geometry of a nanopipette as a simplified truncated hollow cone immersed in a circular bath (Figure S1.1). The model was informed by experimental measurements (scanning electron microscopy micrographs of the nanopipette tip geometry, bulk conductivities and viscosities of the solutions), and only the inner half-cone angle (*θ*), the surface charge of the quartz glass (*σ*) and the diffusion coefficients of the solution in the bath containing 50% PEG 35K 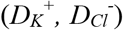 could not be experimentally measured. The inner-half cone angle (*θ* = 7°) was determined parametrically as by solving analytically the resistance of the nanopipette immersed in a 0.1 M KCl solution(S1.2, SI1 for more details). Similarly, the surface charge at the nanopipette quartz wall 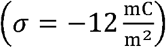 was estimated using the closest fit to the experimental data (Section S1.3, SI1).

In our system, charge is carried by ions migration due to the presence of an electric field (electromigration), concentration gradient (diffusion) and fluid flow (convection)^16^. In 0.1 M KCl, the ion flux generated by electromigration 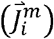 depends on the sum of the diffusion coefficients of ions in solution (S1.4, SI1) which defines the solution conductivity according to the following equation:

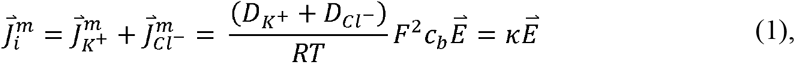

where 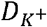 and 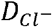 are the diffusion coefficients of potassium and chloride, respectively, *c_b_* the bulk concentration, *F* the Faraday constant, *R* the natural gas constant, *T* the temperature, *κ* the solution conductivity, and 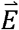 the electric field. In normal conditions (no PEG), the ratio between the diffusion coefficients of the two species is very close to unity 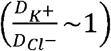, meaning that the contribution of potassium and chloride to the total conductivity *κ* is approximately the same.

Evidence in literature has shown that polyethylene glycol associates with cations in solution^20–23,29^. Zhang et al.^19^ proved experimentally the interaction between cations and PEG, finding that the trapping time of the ion in the polymer chain is highly dependent on the ion radius with longer trapping time for larger radii. These findings are a clear indication that the diffusion properties of cations in solution are affected in the presence of PEG. In the simulations, we considered this effect by assuming an imbalance between the diffusion coefficients of the two ion species in the bath solution. The properties of the 0.1 M KCl electrolyte inside the nanopipette were kept constant as described above.

We performed a parametric study to determine the ratio of the diffusion coefficients by decreasing the contribution of the potassium ion and increasing the one of chloride 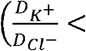 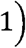 to the total conductivity *κ_PEG_* (Section S1.4, Supporting Information 1) to describe the experimental *i-V* of the nanopipette in the presence of PEG shown in Figure 1b (orange curve). This study revealed that the higher the ratio of diffusion coefficients, the more asymmetric the *i-V* response will be (Figure S1.4, SI1), which supports our hypothesis that the polymer-cation interactions are responsible for the distinctive current response in presence of PEG^18,30^. We obtained the closest fit to the experimental data (orange square points, Figure 1b) by selecting a diffusion coefficient ratio of 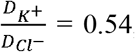, meaning a 35% contribution from the cations and 65% from the anions to the total conductivity of the PEG solution. The simulated currents shown in Figure 1b (orange data points) quantitatively reproduce the experimentally observed *i*-*V* response (orange curve).

It is worth clarifying that all inputted parameters, with or without PEG in the outer solution, were either measured experimentally (electrical conductivity, fluid viscosity and electrolyte concentration) or found in literature (electric permittivity, fluid viscosity) (Table S1.2, SI1). In addition, the nanopipette surface charge and any fluid flow in the system minimally influence the simulated *i*-*V* response in the presence of PEG in the bath solution (Section 1.5 and table S1.3 of Supporting Information 1), thus all modelling results related to PEG presented below were obtained without considering these factors.

### Ion concentrations at the tip region

We then investigated the distribution of ions near the tip of the nanopipette to better understand the effect of PEG in the most sensitive region of our system.

Figure 2 shows the average ion concentration 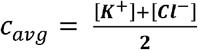 obtained with finite element modelling under two opposite voltages applied (*V* = ±500 mV) in presence (Figure 2a, 2b) and absence (Figure 2c, 2d) of PEG in the external solution (Section S2, SI 1). In the presence of PEG, a pronounced ion depletion is observed for *V* = −500 mV (Figure 2a) while ion enrichment is noticeable when *V* = +500 mV (Figure 2b) with a 20-fold increase in ion concentration compared to when a negative bias is applied. This observation is the origin of the asymmetric current response observed in presence of PEG (Figure 1b). In absence of PEG in the external solution, a slightly higher ion concentration can be observed within the pore region under *V* = −500 mV (Figure 2c) compared to the case with *V* = +500 mV (Figure 2d).

**Figure 2.**
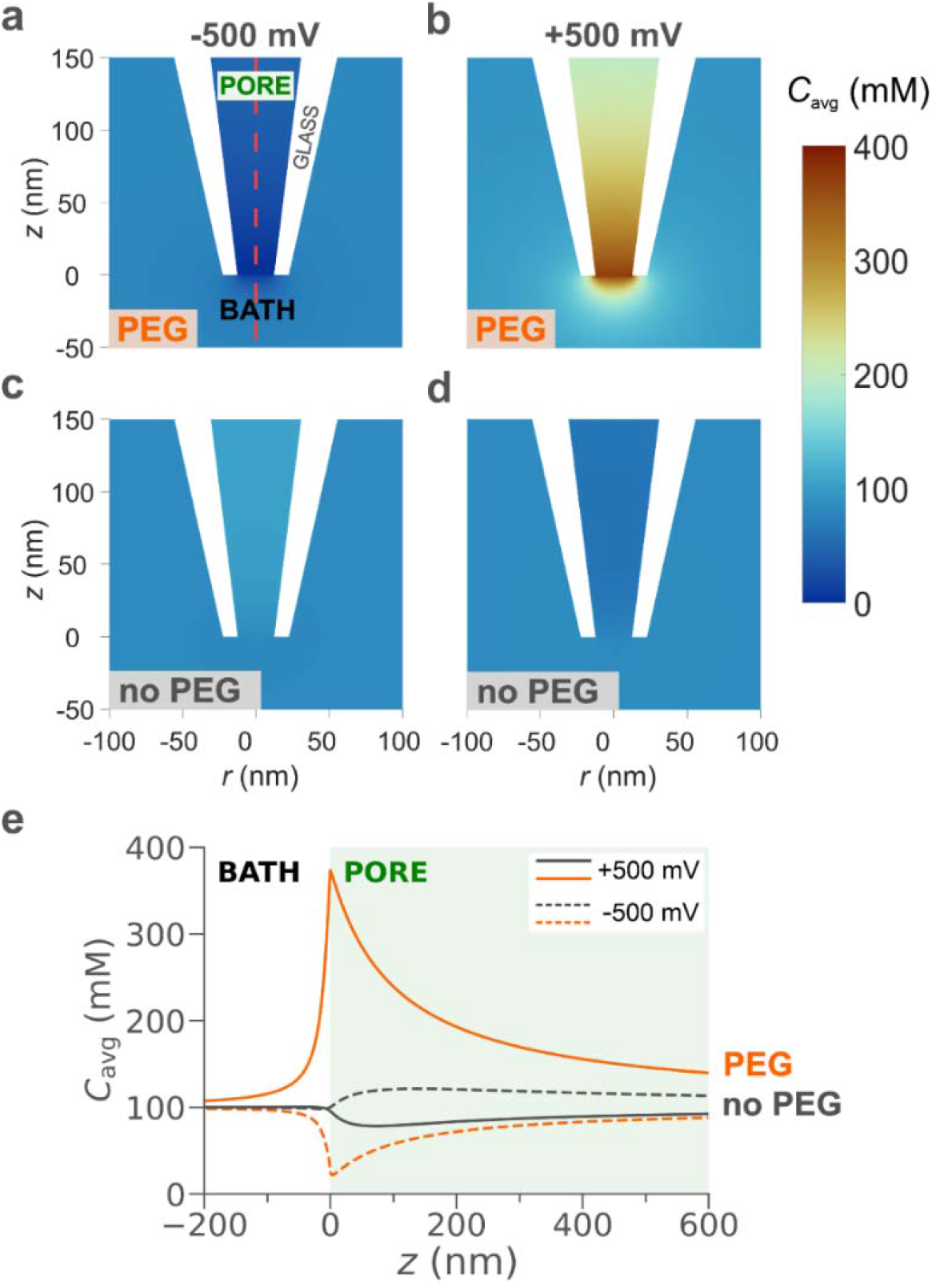
Simulated ion distributions close to the nanopipette tip at ±500 mV in the presence and absence of PEG in the bath solution. Average concentration (*C_avg_* = ½ ([*K*^+^] + [*Cl*^−^])) with (a, b) and without (c, d) PEG in the bath solution for an applied voltage of (a, c) −500mV and (b, d) 500 mV. (e) Average ion concentrations along the nanopipette axis of symmetry (red dashed line in a) in presence (orange) and absence (black) of PEG for negative (dashed curves) and positive (solid curves) bias applied. The diameter of the nanopipette is 25 nm and the internal and external solution is 0.1 M KCl for both PEG and no PEG but in the PEG case, the external solution also contains PEG 35K.

This observation explains the slightly asymmetric curve (ion-current rectification) for the no PEG case (gray curve) shown in Figure 1b^17^.

Figure 2e plots the average ion concentration along the symmetry axis of the pipette (dashed red line, Figure 2a), allowing for quantitative comparison of the simulations. The average concentration for the PEG (orange curve) and no PEG (gray curve) case is plotted for *V* = −500 mV (dashed line) and *V* = +500 mV (solid line). In our reference system, the interface between nanopipette and external solution is positioned at *z* = 0 nm, while *z* > 0 nm correspond to the axis of symmetry inside the nanopipette and *z* < 0 nm to the external solution (Figure S1.1). Interestingly, the maximum ion concentration for *V* = +500 mV in the presence of PEG (orange solid line) is approximately 4 times higher than the corresponding case with no PEG (gray solid line). This observation indicates that the above- bulk conductivity arises from a dramatic increase in the ion concentration in the sensing region of the nanopipette, despite the external solution in the presence of PEG being 9 times less conductive.

Experimentally, a similar increase in conductivity is observed upon the translocation of a single dsDNA molecule in presence of PEG in the bath solution, as shown in Figure 1c, suggesting that the signal amplification is related to the number of ions in the sensing region of the nanopipette. The vast difference in ion concentration between positive and negative bias is similar to the behavior of nanofluidic diodes^31–35^ for ultrashort conical nanopores. In these studies, nanofluidic diodes were developed by introducing a surface charge discontinuity on a nanochannel which forms a junction similar to bipolar semiconductors. In our case, we achieve a similar behavior by introducing an interface where the value for the diffusion coefficient for cations and anions is approximately the same to a region where the diffusion coefficient for the cations is much smaller than the one for anions due to the presence of PEG. This discontinuity not only affects ions distribution but also ion transport, as we describe in the next section.

### Ions transport at the tip region

The origin of the significant differences in ion concentration (*c_avg_*) in the presence of PEG can be understood by a careful analysis of the ion transport 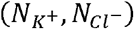 across the interface close to the nanopipette tip aperture, which represents the most sensitive region of our system^36^ (Section S3, Supporting Information 1).

We define “sensing region” as the region between two equipotential lines where a 50% drop of the applied voltage is observed. In case of −500 mV applied, the voltage drop across the sensing region is equal to 250 mV. In presence of PEG and under −500 mV, we found that this region is about 40 nm in length along the z-axis (from z = − 20 nm to z = 20 nm with interface between inner solution and bath solution set at z = 0) (Figures S3.3 and S3.4, SI1). This clearly indicates a highly resistive region positioned at the nanopipette tip which leads to a significant drop in the measured current magnitude, as shown in Figure 1b (orange curves and square points).

In any enclosed volume, the flux of ions through the surface surrounding the volume is equal to the rate of change in the ions number (mass and charge conservation)^37^. The transport rate for each ion species (*N_i_*) was calculated by integrating the total flux of K^+^ and Cl^−^ separately, along the equipotential lines (dashed lines, Figure 3) selected around the nanopipette tip. An extensive description of these calculations is provided in section 3.2 of the Supporting Information. In a nutshell, for 0.1 M KCl, where both ion species have a valence of *z_i_* = 1, the difference between the number of charges (ions) entering and exiting each dashed line over time is proportional to the current.

**Figure 3.**
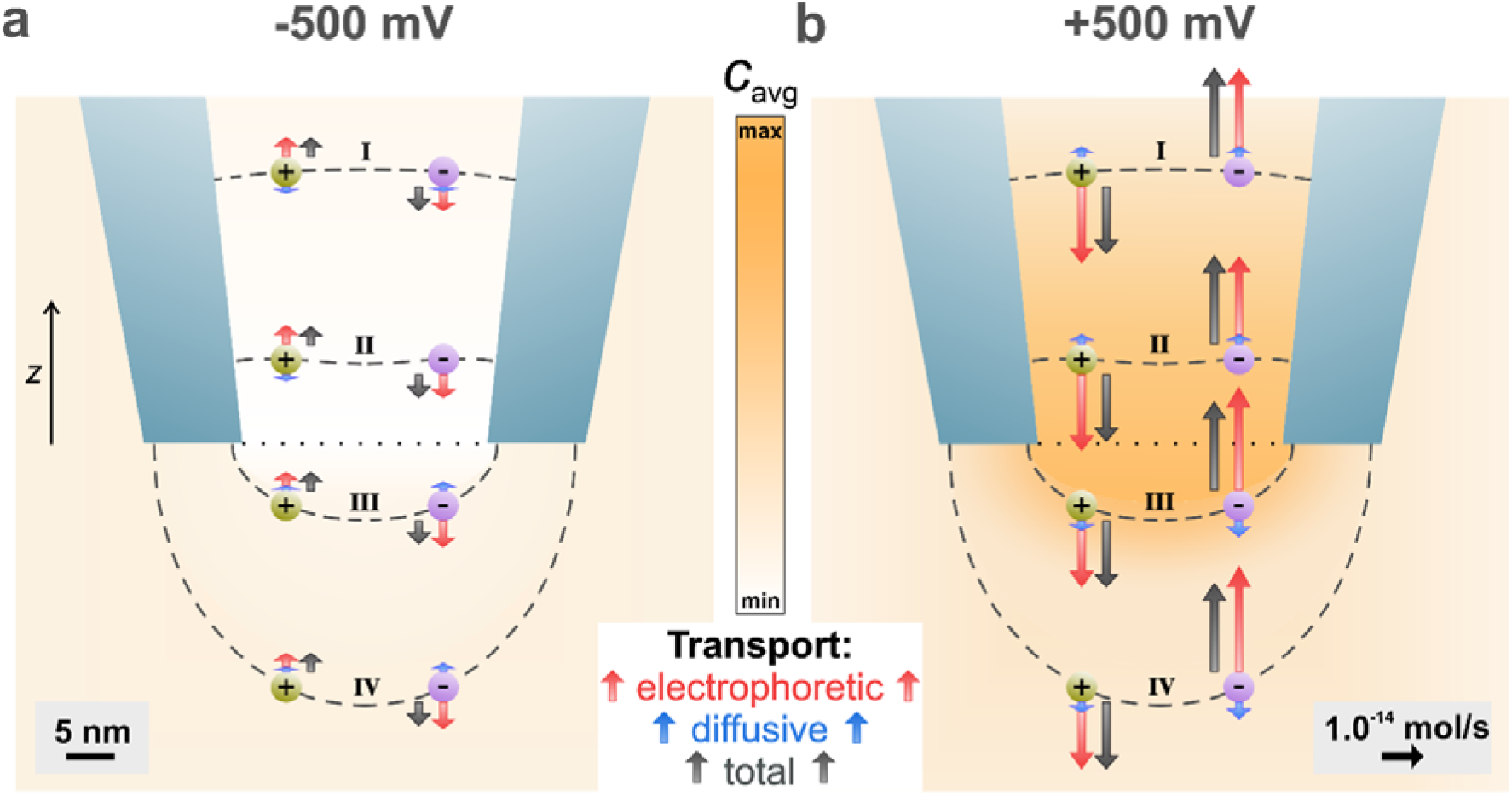
Visualization of the relative contributions of different physical processes to the transport rates of K^+^ and Cl^−^ at ± 500 mV with PEG in the outer solution. The lengths of the arrows represent the magnitude of the total transport rate (gray) across the respective equipotential line (dashed black), which is the sum of electrophoretic (red) and diffusive (blue) contributions. In addition, the arrows being parallel to the z-axis and the ions positions were selected for illustration purposes only. Arrows for negligible diffusive contributions are not shown in the plot for ease of representation. The colour map in the background represents the average ion concentration and the dotted line at the nanopipette aperture the interface between the inner and outer solution.

Since no convection was considered for this simulation, the total ion transport rate (black arrow, Figure 3) can be broken down to two components, the electrophoretic 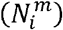 and diffusive 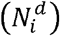 (red and blue arrow, respectively, Figure 3). Figure 3 illustrates all these three components, for both cations (green sphere) and anions (purple sphere), at 4 equipotential lines to highlight the marked difference in ion transport between the inner and outer solution for V = ± 500 mV. The total ion transport rate (black arrows) of each ion species for each applied potential remains constant across the designed dashed lines, verifying that mass and charge is conserved in the system and that the sum of the electrophoretic and diffusive components will always be the same. Based on the polarity of the applied voltage, cations/anions will get attracted/repelled resulting in electrophoretic ion transport either in or out the nanopipette tip (dotted black line, Figure 3). Additionally, any gradients in the ion concentration (color map in background of Figure 3) give rise to diffusive ion transport with both species moving towards (with V = −500 mV) or away the tip interface (with V = 500 mV).

Figure 3 shows that the total ion transport rate at −500 mV is lower than the rate at 500 mV by 75% which is in agreement with the experimental and simulated *i*-*V* responses presented in Figure 1b. It is important to note that the electrophoretic transport dominates diffusion in all cases. In Figure 3a, chloride anions move towards the outer solution, while potassium cations flow towards inside the pore opening. The directionality of transport is exactly opposite in Figure 3b with Cl^−^ moving inside the nanopipette tip and K^+^ travelling outwards. To summarize, when V = 500 mV, there is a larger number of ions flowing across the nanopipette tip aperture over time which results in a higher current magnitude (Table S4.1, SI1) demonstrating that an asymmetric ion mobility is responsible for the observed above-bulk conductivity. In contrast, when V = −500 mV, there is a low number of ions flowing across the nanopipette tip aperture over time resulting in a much lower current magnitude (Table S4.2, SI1) which again is consistent with the experimental data.

### Mechanism of current enhancement upon dsDNA translocation

DNA molecules carry a negative surface charge and form counter-ion clouds when immersed in electrolyte solutions (0.1 M KCl). In standard conditions (no PEG) and under negative potentials (−500 mV), the temporary increase in the current magnitude recorded during dsDNA translocation is due to the additional ions carried by the molecule to the sensing region of the nanopipette which results in a temporary higher ion concentration^4^.

In the presence of PEG, the physics related to the generated current upon dsDNA translocation through the nanopipette aperture is considerably more complex. As previously explained, the nanopipette shows a remarkable ion depletion at the tip region with very few ions transporting through the interface when −500 mV are applied (see ion concentrations in Figure 2a and transport in Figure 3a), while the bath solution is mainly populated by anions with cations transport hindered by intercalation in the PEG molecules. In these conditions, the counter-ion cloud carried by the dsDNA molecule certainly contributes to the temporal increase of the ion concentration, thus the conductivity, of the system. However, this is not sufficient to explain the drastic current enhancement recorded experimentally. In fact, the charge carried by the translocating dsDNA molecule is the same regardless the presence or absence of PEG in the bath solution, thus the increased conductivity should be approximately equal in both cases (see Section S4.1, SI1).

We explored if the mechanical interactions between a dsDNA molecule and the interface between pore could temporarily alter the ion concentrations at the tip region. Briefly, we considered a rectangular protrusion of the domain inside the nanopipette (pore solution) towards the bath domain to get a simplistic model of the interface shift caused by the arrival of DNA, as shown in Figure 4a. We performed a parametric study by varying the size of this protrusion (Δz) from *z* = 0 nm to *z* = −30 nm with 2 nm steps. Figure 4b presents the simulated average ion concentration along the symmetry axis (r = 0 nm) for three different interface displacements (0, 2 and 30 nm). As the interface moves further away from the nanopipette tip opening (z = 0 nm), the number of ions in the nanopipette’s sensing region increases resulting in an enhanced current value. We found that an interface displacement of 16 nm towards is sufficient to cause an increase in the ion current the match the current peak maxima measured experimentally for the translocation of a single 4.8 kbp dsDNA molecule (Section S4.2, SI1). This current enhancement is due to a 33% increase in ion concentration in the nanopipette sensing region (0 < z < 20 nm) caused by this shift in the interface.

**Figure 4.**
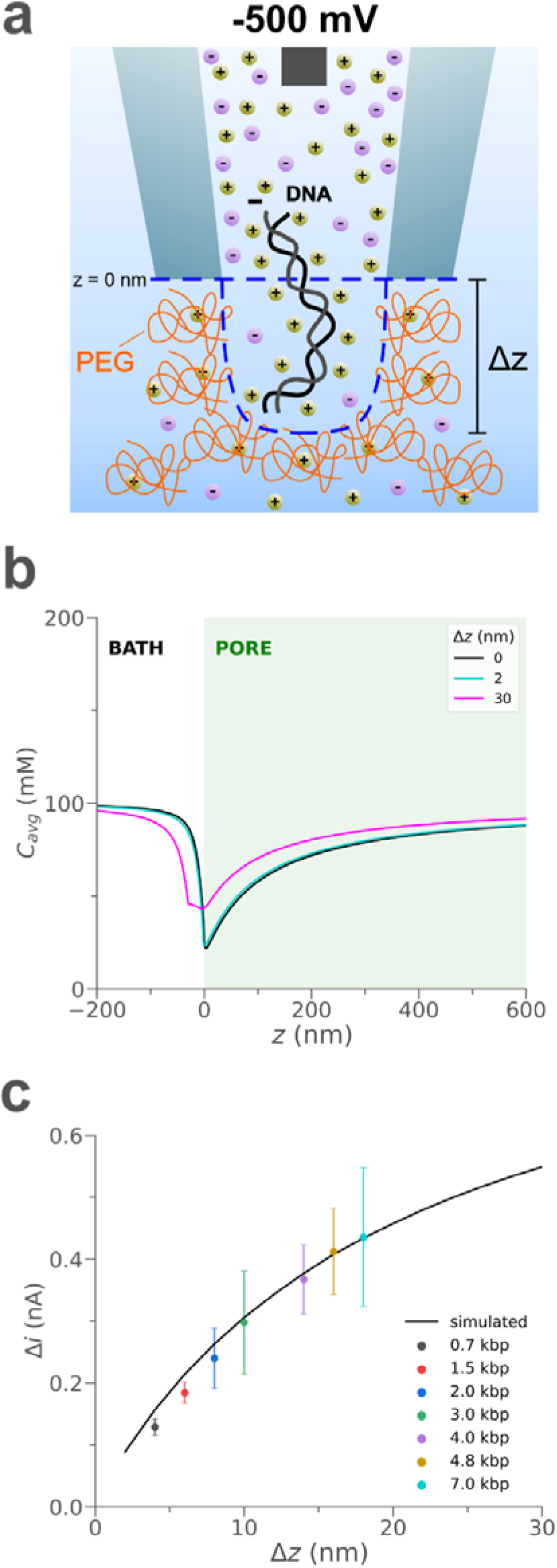
Proposed mechanism of current enhancement upon a dsDNA molecule translocation. (a) The translocation of a dsDNA molecule through the nanopipette causes a temporary displacement of the interface between the pore and bath solution which results in a temporary ion enrichment in the nanopipette tip region (note: the illustrations are not in scale and geometries were chosen for illustration purposes only). (b) Simulated average ion concentration along the axis of symmetry (r = 0 nm) for 0 nm (black), 2 nm (cyan) and 30 nm (magenta) interface displacement. (c) Simulated (black curve) and experimental (coloured points) current peak maxima (Δ*i*) for different interface displacements towards the bath solution and sizes of dsDNA molecules translocating through the nanopipette tip aperture towards the bath, respectively. The error bars represent the standard deviation of the experimental current peak maxima values.

To summarize, we found that the translocation of dsDNA molecules through the pore causes a temporary displacement of the interface, which results in a shift of the ion depleted region towards the bath. The consequence is ion enrichment in the sensing region inside the nanopipette, which results in higher conductivity, thus higher measured currents (Figure 4a). Note that in our simulations, we simplistically assume that the interface between pore and bath solution without DNA is a straight line at z = 0 nm (no mixing, blue dashed line in Figure 4). Using a more sophisticated model for the interface would certainly improve the accuracy of our calculations, but not the level of our understanding of the system.

Based on this mechanism, we expect various dsDNA molecule sizes to have different effects on the translocation current. For instance, longer dsDNA molecules would displace the interface further towards the external solution. To prove this hypothsis, we repeated the same experiment as the one illustrated in Figure 1 using a range of sizes for the analyte (0.7 – 7 kbp) with and without PEG in the outside bath (Figure 4c and Section 4.3, SI1). In PEG, experimental current peak maxima for the translocation of dsDNA molecules with sizes from 0.7 kbp up to 4.8 kbp are in close agreement with the trend obtained from the simulated current values due to interface displacements, as shown in Figure 4c and Table S4.1 in Supporting Information 1. In the no PEG case, not only there is no evident correlation, but the detection is limited to molecules with a minimum size of 4.8 kbp (Figure S4.2c, SI1). These findings confirm our initial hypothesis that the current enhancement in the presence of PEG 35K upon dsDNA translocation cannot be explained only in terms of additional ions carried by the analyte, as recently reported by Lastra et al.^15^ for a system based on a pore’s flux imbalance, but a mechanical interaction between the analyte and PEG molecules at the nanopipette tip opening must be taken into account.

To further support this, we experimentally verify that the voltametric responses and current enhancement caused by PEG disappear when a positive pressure is applied at the back of the nanopipette to push PEG molecules away from the tip opening (Section S4.4, SI1). This result shows that the PEG effect is completely cancelled by disrupting the interface, underpinning the importance of the latter to the experienced current enhancement.

## CONCLUSION

To summarize, we developed a finite element model to improve our understanding of the dramatic current enhancement upon dsDNA molecule translocation through a nanopipette to an external solution containing 50% (w/v) PEG 35K. This system was successfully simulated by assuming asymmetric diffusion coefficients between cations and anions due to the cation binding properties of PEG. We observed that the characteristic *i*-*V* response in the presence of PEG is due to voltage-dependent ion concentrations at the tip region with ion enrichment at positive and ion depletion at negative potentials. A similar behavior was noticed in the asymmetric transport rates for each ion species across the tip orifice, resulting in higher currents at positive applied bias compared to negative. Furthermore, we demonstrated that conventional mechanisms of current enhancement based on additional ions carried by the analyte cannot be fully applied to our system. Hence, we proposed a novel mechanism supported by experimental evidence which relies on mechanical interaction between the translocating analyte and the solutions interface. We proved that such interactions could lead to alteration of the ion distribution at the tip orifice which can result into temporary current increases. We expect that this work can provide a new paradigm in nanopore sensing, where the alteration of the ion transport properties of the solution can be harnessed to provide enhanced signal to noise allowing for the biochemical and structural analysis of proteins and other biomolecules.

## MATERIALS AND METHODS

### Nanopipette fabrication

Quartz capillaries of 1.0 mm outer diameter and 0.5 mm inner diameter (QF100-50-7.5; Sutter Instrument) were used to fabricate the nanopipette using the SU-P2000 laser puller (World Precision Instruments). A two-line protocol was used, line 1: HEAT 750/FIL 4/VEL 30/DEL 150/PUL 80, followed by line 2: HEAT 625/FIL 3/VEL 40/DEL 135/PUL 150. The pulling protocol is instrument specific and there is variation between different SU-P2000 pullers.

### Electrolyte bath preparation

To generate 10 ml of the 50% (w/v) poly(ethylene) glycol (PEG 35K) (Sigma Aldrich 94646), 1 ml of 1 M KCl solution, 4 ml of ddH2O and 5 g of PEG 35K were mixed inside a tube. The tube was then left inside a 70°C incubator for 2 hours followed by overnight incubation at 37°C. The tubes were then left on bench for 4 hours to reach the room temperature prior to use. All electrolytes were stored at room temperature.

### Ion current trace recording

The nanopipettes were all filled with 0.1 ng/μl dsDNA diluted in 0.1 M KCl (P/4240/60; Fisher Scientific) and fitted with a Ag/AgCl working electrode. The nanopipettes were immersed into the electrolyte bath with a Ag/AgCl reference electrode containing or not containing Polyethylene Glycol 35K. The ionic current trace was recorded using a MultiClamp 700B patch-clamp amplifier (Molecular Devices) in voltage-clamp mode. The signal was filtered using low-pass filter at 10 kHz and digitized with a Digidata 1550B at a 100 kHz (10 μs bandwidth) sampling rate and recorded using the software pClamp 10 (Molecular Devices).

### Finite Element Modelling

Finite element simulations were performed with COMSOL Multiphysics 5.6 (COMSOL Inc.). Details for the boundary conditions and meshing are provided in Supporting Information 1 and 2.

## ACKNOWLEDGEMENTS

We thank Prof Joshua B. Edel (Imperial College London) for generously providing the MATLAB script used for event analysis in this study. We thank Prof Aleksei Aksimentiev (University of Illinois, Urbana Champaign) for illuminating discussions. We thank Dr Nataricha Phisarnchananan (University of Leeds) for performing the viscosity measurement of the electrolyte.

