## Supporting Information for "Asymmetric Ion Mobility and Interface Displacement Drive the Signal Enhancement in a polymer-electrolyte nanopore"

**Contents**

**S1	Simulation overview and verification**

*S1.1 Finite element model overview*

*S1.2 Comparing simulated and analytical nanopipette resistance*

*S1.3 Fitting simulation to experimental data without 50% PEG in the bath solution*

*S1.4 Fitting simulation to experimental data with 50% PEG in the bath solution*

*S1.5 Effect of wall surface charge and fluid flow on finite element model*

*S1.6 Changes in experimental i-V response based on external solution composition*

**S2 Ion concentrations at the tip region**

**S3 Ions transport at the tip region**

*S3.1 Updated finite element model overview*

*S3.2 Calculating the transport rates of each ion species at the designed boundaries*

*S3.3 Defining the “sensing region” based on the electric potential distribution*

*S3.4 Transport rates of each ion species in “sensing region” with PEG in the bath*

**S4	Mechanism of current enhancement upon dsDNA translocation**

*S4.1 Estimating the number of ions carried by single dsDNA in the “sensing region”*

*S4.2 Model for interface displacement towards outside bath due to dsDNA translocation*

*S4.3 Effect of dsDNA size on translocation current*

*S4.4 Experimental and simulated effect of applied pressure to the nanopipette*

**References**

**S1	Simulation overview and verification**

*S1.1 Finite element model overview*

In this work, we designed a finite element model, based on a simplified geometry of a nanopipette tip (truncated cone) immersed in an electrolyte solution containing 50% (w/v) PEG, to solve the coupled Nernst-Planck, Poisson and Navier-Stokes equations and understand the physical mechanisms involved. By allocating the appropriate boundary conditions on this two-dimensional (2D) axisymmetric model (Figure S1.1), we determined the ion concentration, voltage and velocity distributions in the fluid. For the solution of these equations, we assumed steady-state conditions $\left( \nabla{\overset{⃑}{J}}_{i}=0 \right)$ and incompressible fluid flow $\left( \nabla\overset{⃑}{u}=0 \right)$.

The Nernst-Planck equation (“*Transport of Diluted Species*, *chds*”) describes the ion transport properties based on the diffusive, electrophoretic and convective flux, as follows^1^:

$${\overset{⃑}{J}}_{i}=-D_{i}\nabla c_{i}+\frac{z_{i}F}{RT}D_{i}c_{i}\nabla V-c_{i}\overset{⃑}{u}$$

(S1.1),

where ${\overset{⃑}{J}}_{i}$: ion flux, *D_i_*: diffusion coefficient of ion, *c_i_*: ion concentration, *z_i_*: ion valence number, *F*: Faraday constant, *R*: universal gas constant, *T*: temperature, *V*: voltage and $\overset{⃑}{u}$: fluid velocity.

The Poisson equation (“*Electrostatics*, *es*”) describes how the electric potential is related to the ion concentration:

$$\nabla^{2}V=-\frac{F}{\varepsilon}*\sum_{i} z_{i}c_{i}$$

(S1.2),

where *ε*: medium electric permittivity.

The Navier-Stokes equation (“*Laminar Flow*, *spf*”) describes the flow distribution:

$$\overset{⃑}{u}\nabla\overset{⃑}{u}=\frac{1}{\rho}\left( -\nabla p+\eta\nabla^{2}\overset{⃑}{u}-F\left( \sum_{i} z_{i}c_{i} \right)\nabla V \right)$$

(S1.3),

where *ρ*: medium density, *η*: medium viscosity and *p*: pressure.

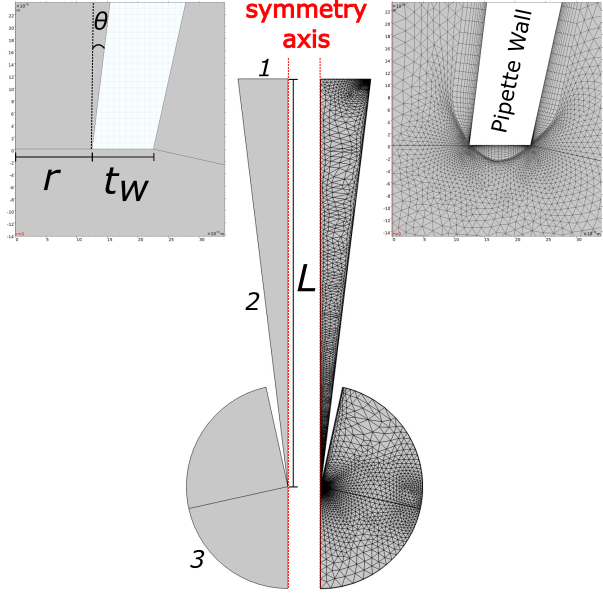

**Figure S1.1** (Left) Zoomed-in view of nanopipette tip geometry with the pore tip radius (r), inner half-cone angle (θ) and quartz wall thickness (t_w_) labelled. (Middle) 2D axisymmetric finite element model geometry of a simplified nanopipette (truncated cone) immersed in an electrolyte solution, including the symmetry axis (red dashed lines), important boundaries (1, 2, 3), pore length (L) and final mesh. (Right) Zoomed-in view of meshed geometry of nanopipette tip opening.

Figure S1.1 illustrates the two-dimensional axisymmetric geometry of the finite element model, including three key boundaries for solving equations S1.1, S1.2, S1.3 and the applied mesh. The nanopipette tip aperture (*r*) is 12.5 nm, quartz wall thickness (*t_w_*) is 10 nm, inner half-cone angle (*θ*) is 7° and the pore length (*L*) is 50 μm. The top boundary (1) represents the bottom surface of the Ag/AgCl electrode immersed in the nanopipette where voltage is applied. Boundary 2 is quartz glass walls allocated with a surface charge (*σ*) equal to -12 mC/m^2^, and the semi-circular boundary at the bottom (3) represents the surface of the ground electrode (Ag/AgCl) in the external solution with 50% (w/v) PEG 35K. Tables S1.1 and S1.2 include the main boundary conditions for the equations explained above and important parameters for the numerical simulations obtained either experimentally or analytically (for more information, see sections S1.2, S1.3 and S1.4) The final mesh contains 25,684 elements with an average mesh element quality of 81.4%. Further details regarding all the steps required to design the geometry of this model, apply the same boundary conditions and mesh are provided in Supporting Information II (COMSOL model report).

**Table S1.1** Key boundary conditions applied in finite element model (no PEG and PEG)

| **Boundary** | **Nernst-Planck** | **Poisson** | **Navier-Stokes** |
| --- | --- | --- | --- |
| Bulk solution (1) | $c=c_{b}=100 mM$ | $V=V_{app}$ | $p=491.24 mPa$ |
| Nanopipette walls (2) | $\hat{n}\cdot{\overset{⃑}{J}}_{i}=0$ | $\sigma=\hat{n}\cdot\epsilon\nabla V$ | $\overset{⃑}{u}=0$ |
| Bath solution (3) | $c=c_{bath}=100 mM$ | $V=0$ | $p=0 Pa$ |

**Table S1.2** Key numerical parameters applied in finite element model (no PEG and PEG)

| **Parameter** | **No PEG** | **PEG** | **Type** |
| --- | --- | --- | --- |
| $D_{K^{+}}$ (m^2^/s) | $1.7*{10}^{-9}$ | $1.4*{10}^{-10}$ | *best fit to model* |
| $D_{Cl^{-}}$ (m^2^/s) | $1.8*{10}^{-9}$ | $2.6*{10}^{-10}$ | *best fit to model* |
| $\kappa$ (S/m) | $1.299$ | $0.151$ | *experimental* |
| $\eta$ (Pa*s) | $9*{10}^{-4}$ | $8.73$ | *experimental* |
| $\sigma$ (mC/m^2^) | $-12$ | $0$ | *best fit to model* |

*S1.2 Comparison of the simulated vs analytically calculated nanopipette resistance*

The resistance of a conical nanopore filled with and immersed in the same electrolyte, which depends on the tip aperture radius (*r*), length of the pore (*L*), electrical conductivity of solution (*κ*) and inner half-cone angle (*θ*), is described by the following equation:

$$R_{p}=\frac{L}{\pi\kappa r\left( r+L*tan\left( \theta\right) \right)}+\frac{1}{4\kappa r}$$

(S1.4),

which for $L*tan\left( \theta\right)\gg r$, becomes ^2^:

$$R_{p}=\frac{1}{\kappa r}*\left( \frac{1}{\pi\tan\left( \theta\right)}+\frac{1}{4} \right)$$

(S1.5).

A patch-clamp amplifier (MultiClamp 700B, Molecular Devices) was used to measure the resistance of a nanopipette (*r* = 12.5 nm), filled with and immersed in a solution of 0.1 M KCl (*κ* = 1.299 S/m), which was equal to 175 MΩ. By inputting this value in Equation S1.5 and solving based on *θ*, the inner half-cone angle of the nanopipette model is approximately 7°. It is worth noting that Equation S1.4 provides the same result for the conical pore (*L* = 50 μm) designed in the model. The resulting simulated *i*-*V* curve obtained for the conical geometry described above, but without considering any surface charge on the wall boundaries (*σ* = 10^-6^ C/m^2^) is ohmic with a slope corresponding to 175 MΩ, as shown in Figure S1.2. The fact that there is no difference between the simulated and analytically solved resistance of this system, is the first step towards validating the finite element simulation we developed in this work.

*S1.3 Fitting simulation to experimental data in 0.1 MK KCl*

Once the simulation verified the ohmic response of the system, we attempted fitting the experimental data obtain in 0.1M KCl (Figure 1b, grey curve), by adding only a negative surface charge to the boundaries representing the quartz nanopipette walls. Based on values found in the currently available literature^2,3^, the ion current rectification (ICR) of the system was studied for surface charge values ranging from -8 to -24 mC/m^2^. Figure S1.3 illustrates the simulated *i*-*V* curves for all these values with σ = -12 mC/m^2^ providing the closest match to the experimentally measured *i*-*V* curve.

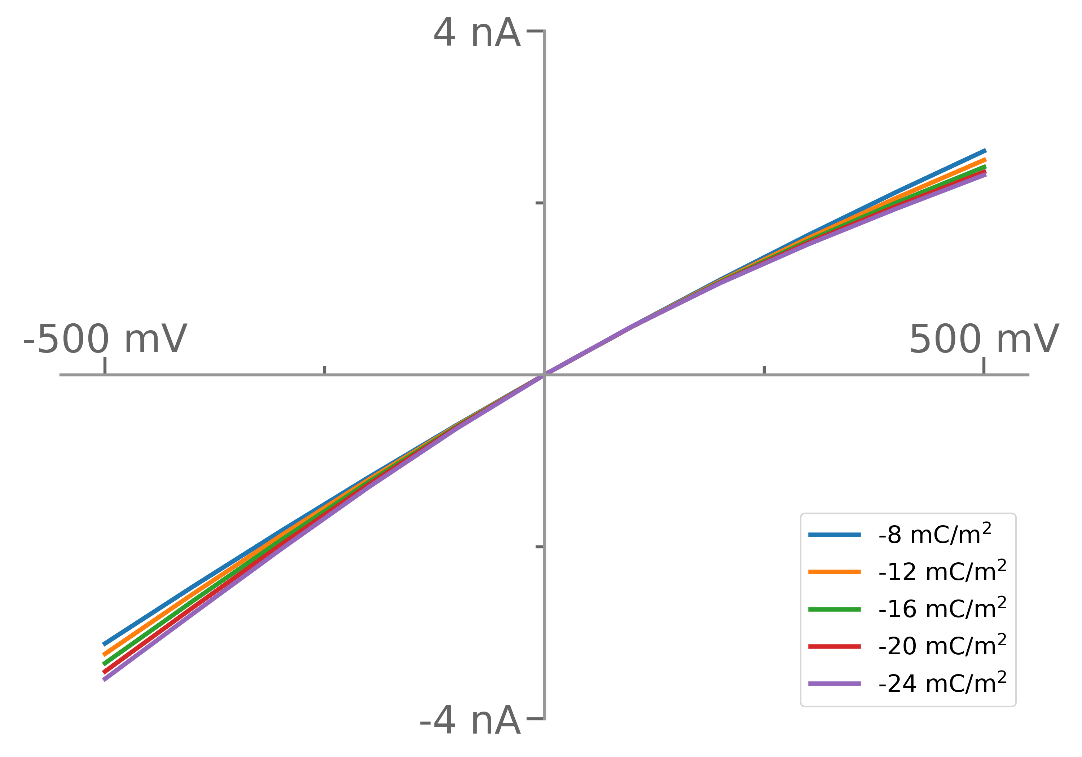

**Figure S1.3** Simulated voltammograms when no PEG is added in the external solution, for a range of applied surface charges on the nanopipette walls. The value -12 mC/m^2^ (orange) provided the closest match to the experimental i-V.

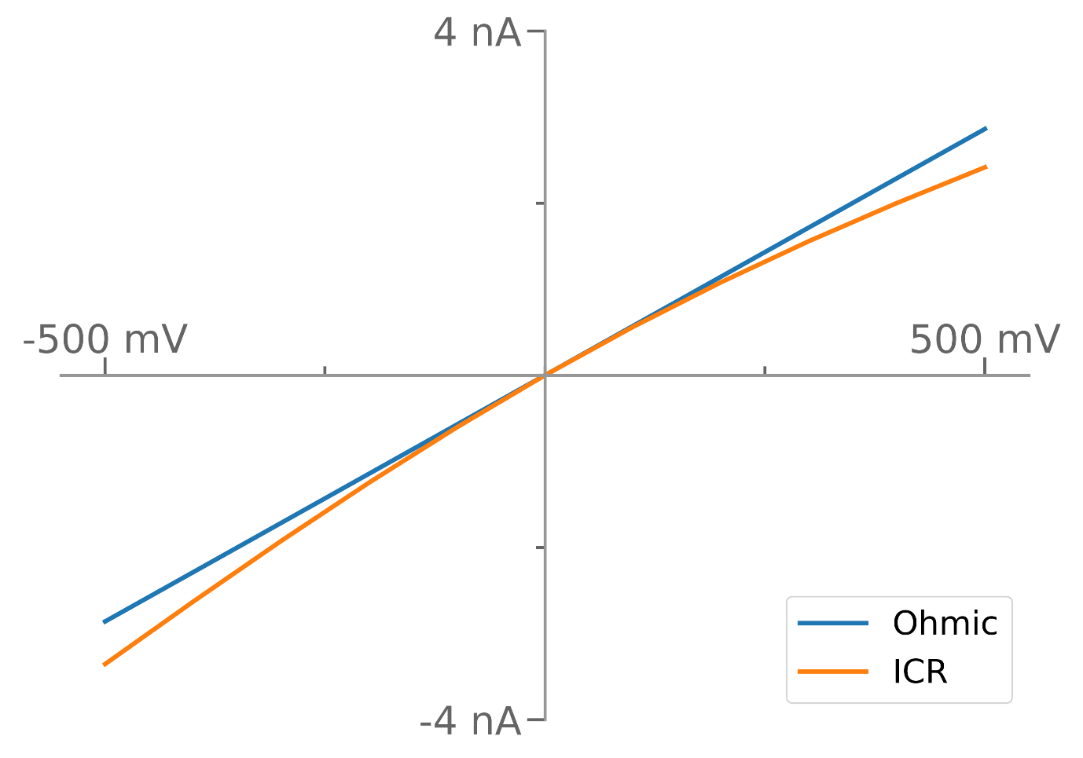

**Figure S1.2** Simulated voltammograms in 0.1M KCl, verifying ohmic i-V response (blue) when no surface charge (σ = 0 mC/m^2^) is applied on the nanopipette walls and negative ion current rectification (ICR, orange) when σ = -12 mC/m^2^.

*S1.4 Fitting simulation to experimental data in 0.1M KCl with 50% PEG*

In any electrolyte solution, the ion flux generated by electromigration $\left( {\overset{⃑}{J}}_{i}^{m} \right)$ is described by Equation S1.6:

$${\overset{⃑}{J}}_{i}^{m}=-\frac{z_{i}F}{RT}D_{i}c_{i}\overset{⃑}{E}$$

(S1.6),

where $\overset{⃑}{E}$**:** the electric field^4^. In our system, where $c_{b}=$ 0.1 M KCl, the total electrophoretic flux is equal to the sum of fluxes contributed by the cations $\left( {\overset{⃑}{J}}_{K^{+}}^{m} \right)$ and anions $\left( {\overset{⃑}{J}}_{{Cl}^{-}}^{m} \right)$, as shown below:

$${\overset{⃑}{J}}^{m}={\overset{⃑}{J}}_{K^{+}}^{m}+{\overset{⃑}{J}}_{{Cl}^{-}}^{m}= \frac{\left( D_{K^{+}}+D_{Cl^{-}} \right)}{RT}F^{2}c_{b}\overset{⃑}{E}=\kappa\overset{⃑}{E}$$

(S1.7).

In dilute aqueous solutions, the bulk diffusion coefficients of K^+^ and Cl^-^, are similar with values of $D_{K^{+}}=1.957*{10}^{-9} \left( \frac{m^{2}}{s} \right)$ and $D_{{Cl}^{-}}=2.032*{10}^{-9}\left( \frac{m^{2}}{s} \right)$, respectively^5^. By adding these two values, we found that potassium cations contribute 49% to the total conductivity while chloride anions contribute the remaining 51%. Based on this and Equation S1.7, we defined the diffusion coefficient for each ion species through the experimentally measured conductivity $\left( \kappa_{PEG}=0.15\frac{S}{m}, \kappa_{no PEG}=1.3\frac{S}{m} \right)$ as follows:

$$D_{K^{+}}=0.49*\frac{RT}{F^{2}c_{b}}*\kappa$$

(S1.8)

$$D_{{Cl}^{-}}=0.51*\frac{RT}{F^{2}c_{b}}*\kappa$$

(S1.9)

When PEG is added to the external bath, a 49/51 contribution of the ionic species to the total conductivity cannot explain the anomalous *i*-*V* response. We therefore considered the published evidence of the cation-binding properties of PEG ^6^. PEG association with cations in solution was defined by considering an imbalance between the diffusion coefficients of the two species $\left( \frac{D_{K^{+}}}{D_{Cl^{-}}}<1 \right)$. A parametric study was performed by increasing the chloride contribution and decreasing the potassium contribution to the total conductivity until the best fit to the experimental *i*-*V* was obtained (Figure S1.4). The best fit was found for $\frac{D_{K^{+}}}{D_{Cl^{-}}}=0.54$, meaning that the contribution of potassium to the total conductivity in presence of PEG is 35% while chloride contributes to the remaining 65%, according to the following equations:

$$D_{K^{+}}^{PEG}=0.35*\frac{RT}{F^{2}c_{b}}*\kappa_{PEG}$$

(S1.10)

$$D_{{Cl}^{-}}^{PEG}=0.65*\frac{RT}{F^{2}c_{b}}*\kappa_{PEG}$$

(S1.11)

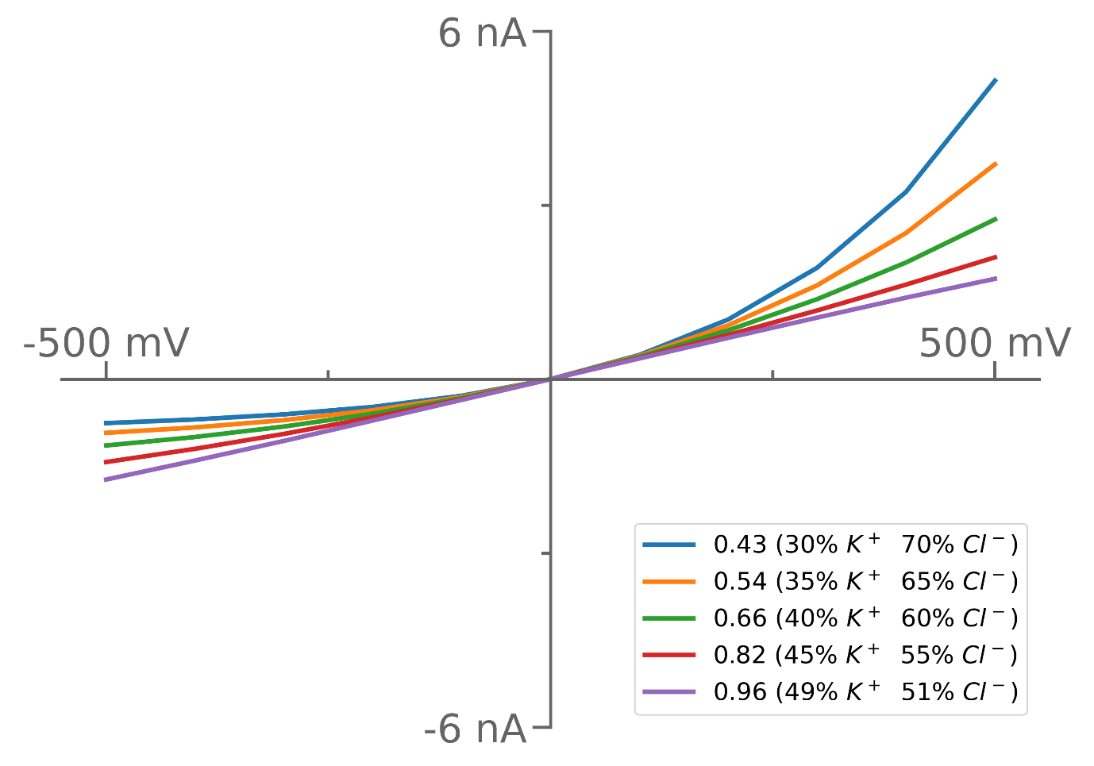

**Figure S1.4** Simulated voltammograms for different ratios between the diffusion coefficient of K^+^ and Cl^-^. The orange curve (0.54) was found to be the best fit to the experimental i-V curve.

*S1.5 Effect of the nanopipette surface charge and fluid flow*

Once we obtained all the parameters for the PEG bath (external domain) of the finite element model, we investigated how the surface charge applied on the nanopipette wall boundaries and any fluid flow (*spf*) in the system affected the solution of the simulation. As Table S1.3 presents, after solving the model by activating either the surface charge, the laminar flow, both or none, there were no significant changes in the calculated current for the applied voltage range [-500 mV, +500 mV]. As a result, we decided to deactivate any surface charge on the nanopipette wall boundaries or laminar flow in the system to simplify the simulation solution and to help us illustrating better the concentration of all ion species.

**Table S1.3** Simulated voltammograms for the finite element model with PEG in the external solution when a combination of the surface charge applied on the nanopipette wall boundaries and any laminar flow in the system are activated/deactivated.

| *V* (mV) | *i* (nA) | | | |
| --- | --- | --- | --- | --- |
|  | $\sigma= 0 mC/m^{2}$  No laminar flow | $\sigma= 0 mC/m^{2}$  Laminar flow | $\sigma=-12 mC/m^{2}$  No laminar flow | $\sigma=-12 mC/m^{2}$  Laminar flow |
| -500 | -0.93 | -0.93 | -0.99 | -0.99 |
| -400 | -0.93 | -0.83 | -0.88 | -0.88 |
| -300 | -0.70 | -0.70 | -0.73 | -0.73 |
| -200 | -0.53 | -0.53 | -0.55 | -0.55 |
| -100 | -0.31 | -0.30 | -0.31 | -0.31 |
| 0 | 0 | 0 | 0 | 0 |
| 100 | 0.40 | 0.40 | 0.40 | 0.40 |
| 200 | 0.93 | 0.93 | 0.92 | 0.92 |
| 300 | 1.62 | 1.62 | 1.60 | 1.60 |
| 400 | 2.52 | 2.52 | 2.47 | 2.48 |
| 500 | 3.70 | 3.71 | 3.63 | 3.64 |

*S1.6 Effect of viscosity*

To investigate whether voltammograms in PEG can be replicated using a solution with similar viscosity, we experimentally tested the *i*-*V­* responses of a nanopipette filled with 0.1 M KCl when immersed in three different solutions composed of 0.1 M KCl, 0.1 M KCl with 50% (w/v) PEG 35K and 0.1 M KCl with 50% (v/v) glycerol. Figure S1.6 proves that the PEG-related *i*-*V* could not be achieved by adding glycerol in the external 0.1 M KCl solution.

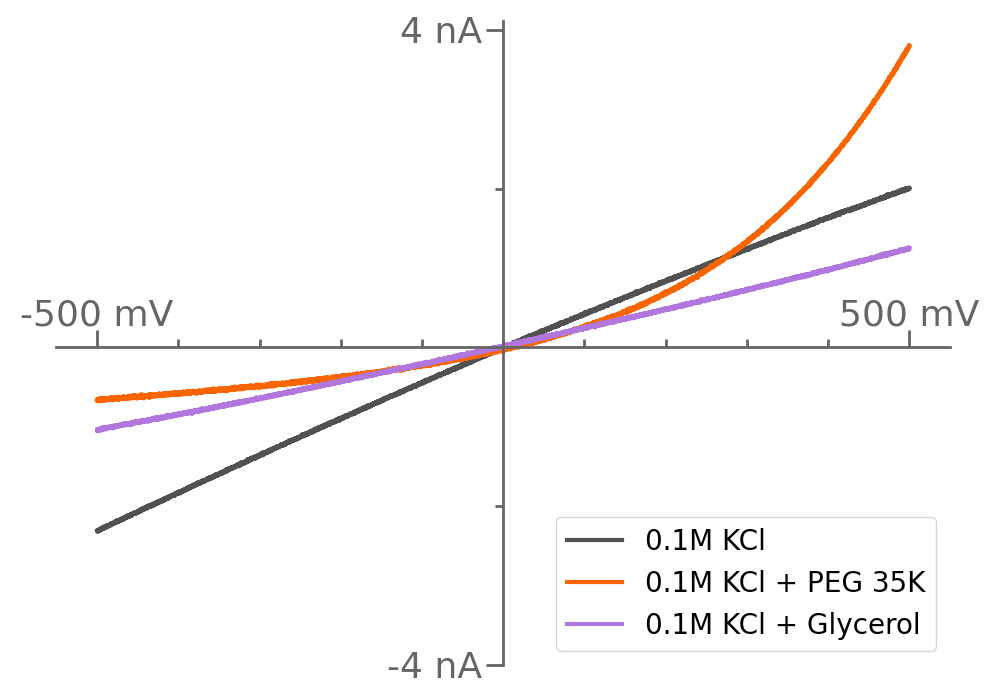

**Figure S1.6** Experimental voltammograms of a nanopipette filled with 0.1 M KCl and immersed in either 0.1 M KCl (grey), 0.1 M KCl with 50% (w/v) PEG 35K (orange) and 0.1 M KCl with 50% (v/v) glycerol (pink).

*S1.7 Analytical estimation of the osmotic pressure in the system*

We analytically estimated the osmotic pressure based on the following equation^7^:

$$p_{osmotic}=icRT$$

(S1.12),

where *i*: van ’t Hoff factor. PEG does not dissociate when dissolved in water, so $i=1$. In the case of 50% PEG 35K being dissolved, by considering a volume $V=0.01 L$, the mass of PEG is $m=5$ g. Given the molecular weight of PEG 35K (MW = 35,000 g/mol), the concentration of PEG becomes $c=0.014 \left( \frac{mol}{L} \right)$. Given these numbers, the estimated osmotic pressure is:

$$p_{osmotic}=36 kPa$$

(S1.13)

Figure S1.7 shows *i*-*V* curves obtained for an osmotic pressure of 0 kPa and 36 kPa with no difference between them.

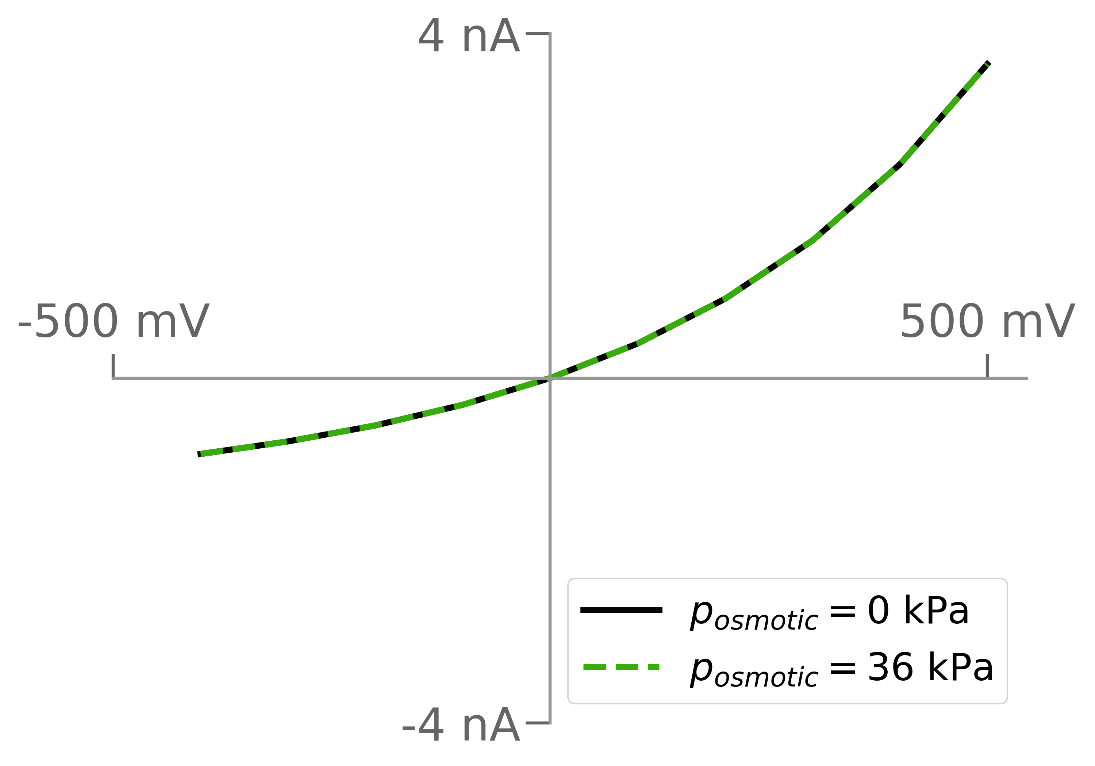

**Figure S1.7** Analytically obtained voltammograms for an osmotic pressure equal to 0 kPa (black line) and 36 kPa (green dashed line).

**S2 Ion concentrations at the tip region**

Figure S2.1 illustrates the cation (K^+^) and anion (Cl^-^) concentrations along a portion of the axis of symmetry (red dashed line, Figure S1.1) when PEG is present (orange) or absent (black) in the external solution under an applied voltage of -500 mV (dashed line) and +500 mV (solid line). Both ion concentrations tend to reach the value of the initially applied concentration (100 mM) for z < -200 nm and z > 600 nm.

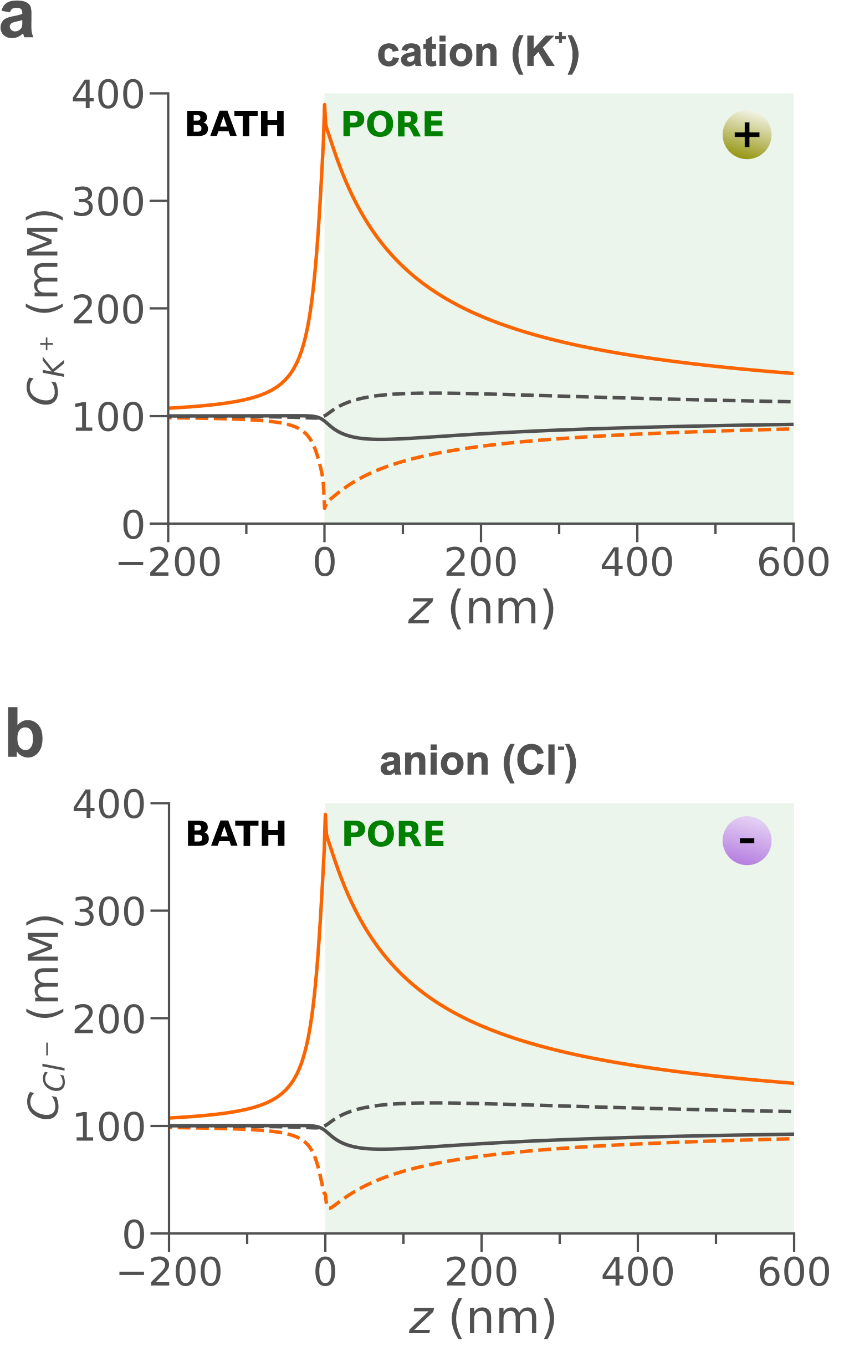

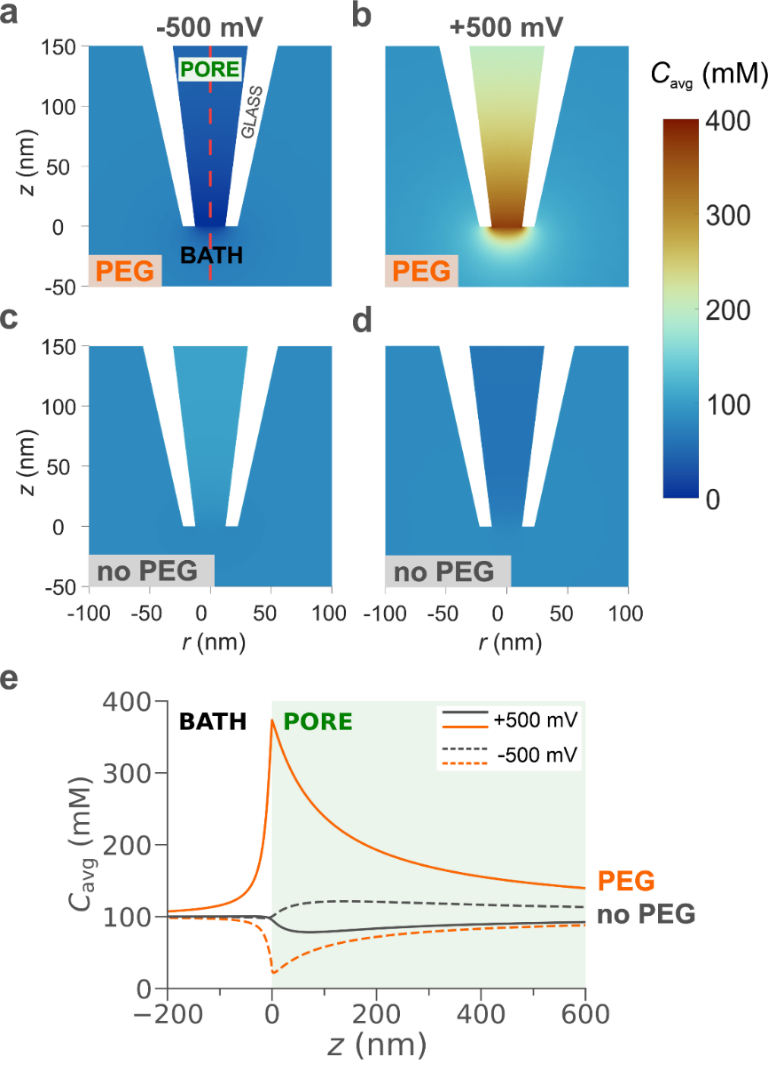

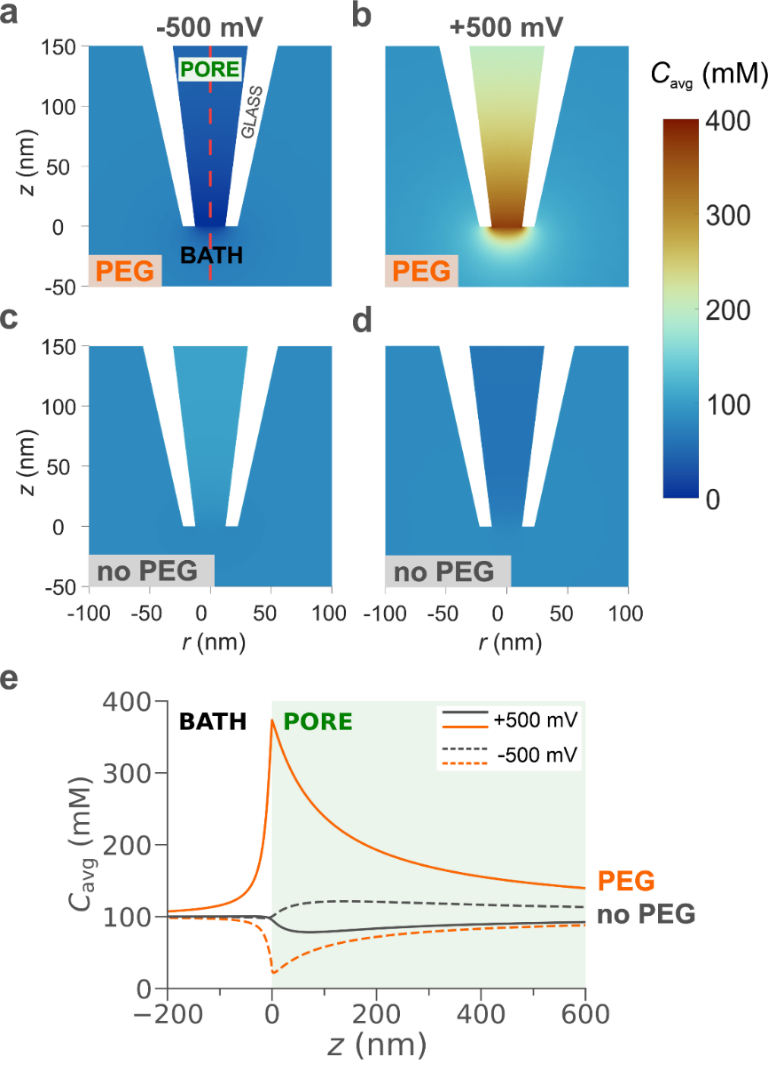

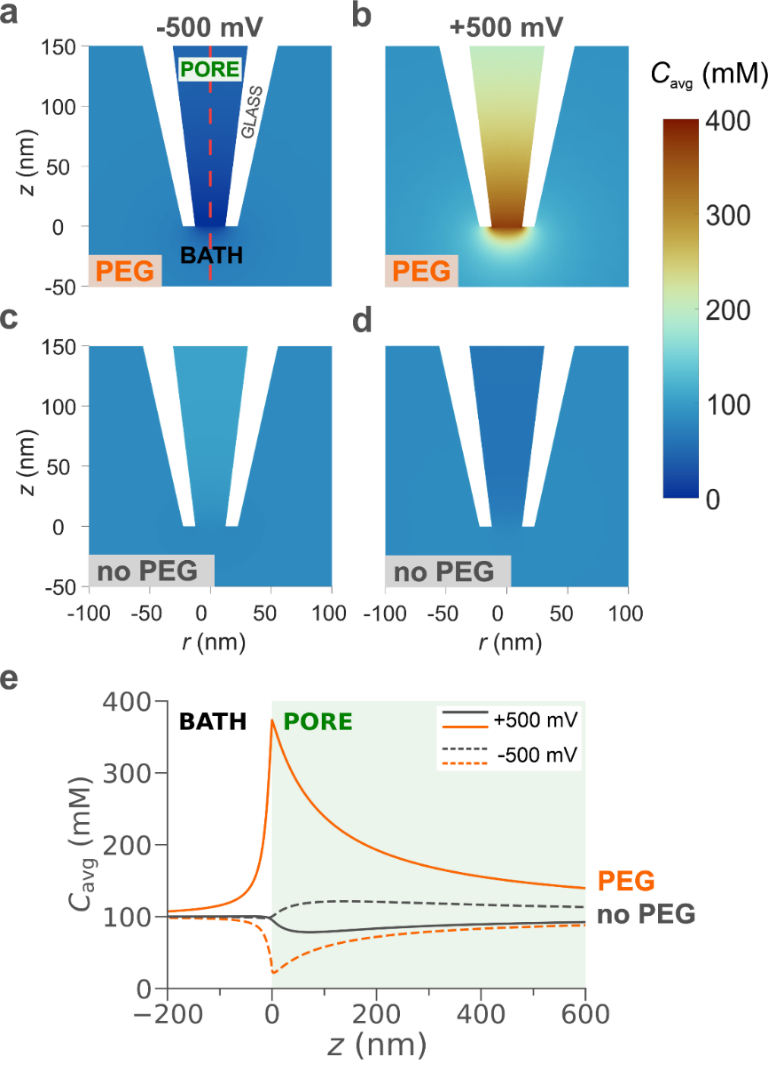

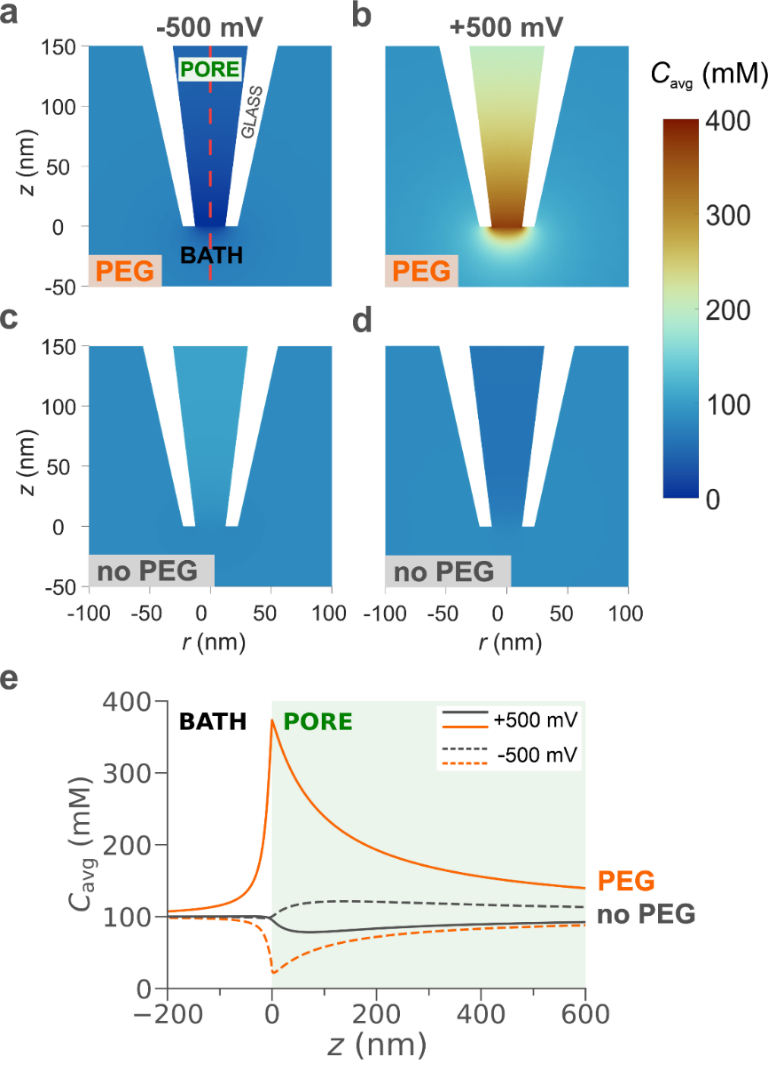

**Figure S2.1** a) Cation (K^+^) and b) anion (Cl^-^) concentrations along 800 nm of the nanopipette axis of symmetry (red dashed line in Figure S1.1) in presence (orange) and absence (black) of PEG for -500 mV (dashed curves) and +500 mV (solid curves). The diameter of the nanopipette is 25 nm and the internal and external solution is 0.1 M KCl for both PEG and no PEG, but in the PEG case, the external solution also contains PEG 35K.

**S3	Ions transport at the tip region**

*S3.1 Boundaries for the calculation of ion transport*

The geometry design analysed in S1.1 and also in Supporting Information II (COMSOL model report), was slightly modified by adding semi-circular and semi-elliptical boundaries around the nanopipette tip opening which could then be used as curves to calculate the integral of the normal ion fluxes, and hence the transport rates of each ion species. Initially, we designed a semi-circle with radius 1.5 μm and centre coordinates (0, 0 nm), which was divided into smaller ones by adding 11 layers, as shown in Figure S3.1a. Then, a semi-ellipse with major semi-axis of 22.5 nm, minor semi-axis of 10 nm and centre at (0, 0 nm) was also added in the geometry, which was also split into 10 smaller ones by reducing the minor semi-axis with a step of 1 nm, except the last which had a step size of 0.7 nm (Figure S3.1b). All these correspond to the simulation where PEG is present in the outer bath.

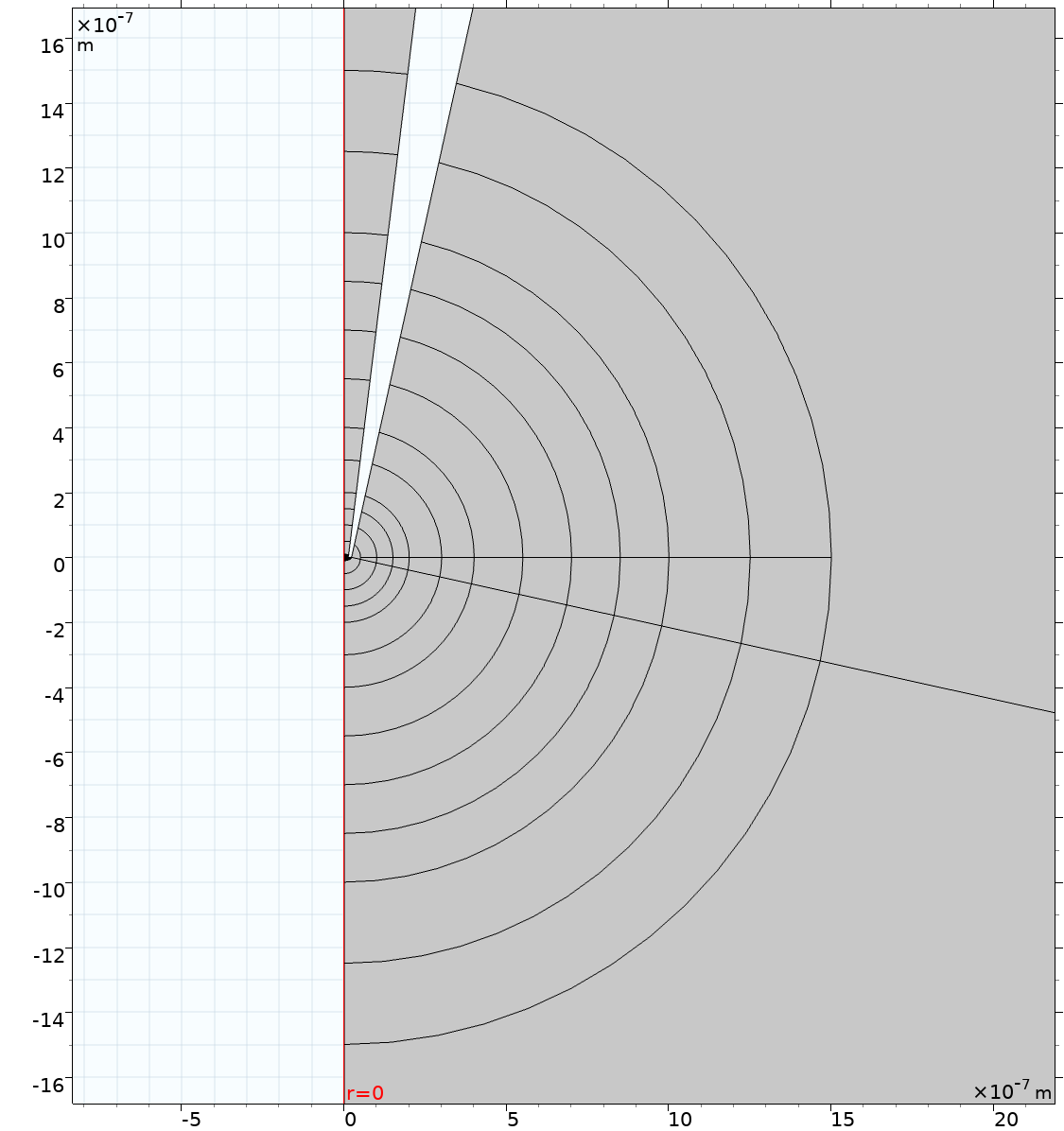

a)

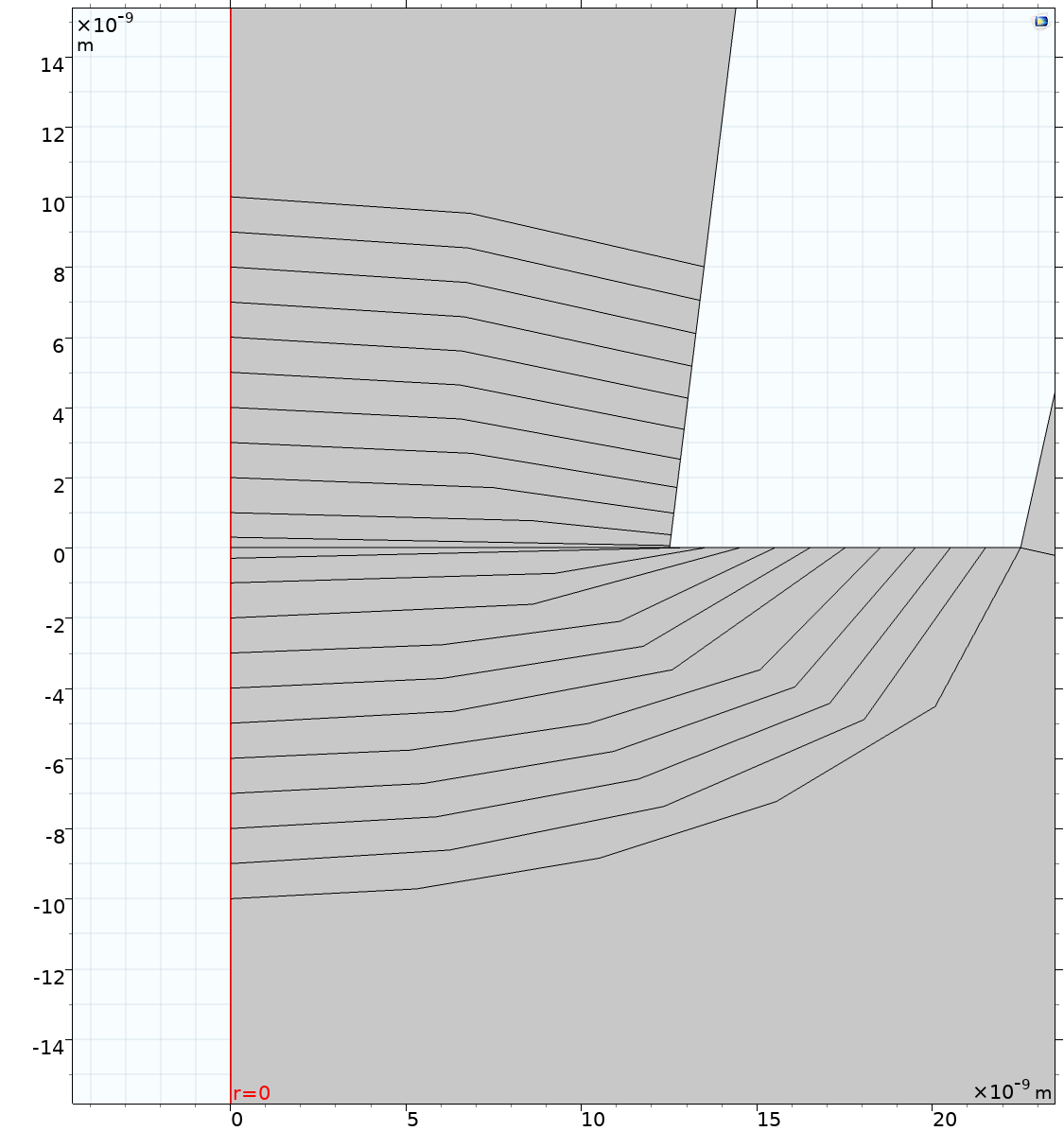

b)

**Figure S3.1** COMSOL zoomed-in geometry plots showing a) semi-circular boundaries inside and outside the nanopipette pore with common centre at (0 nm, 0 nm) and radii ranging from 50 nm to 1.5 μm and b) semi-elliptical boundaries inside and outside the nanopipette pore with common centre at (0 nm, 0 nm) and minor semi-axis ranging from 0.3 nm to 10 nm.

*S3.2 Calculating the transport rates of each ion species at the designed boundaries*

As previously explained, since convection (*Laminar Flow*) did not influence these simulations, the total ion flux $\left( {\overset{⃑}{J}}_{i} \right)$ is only composed of the electromigrative and diffusive ion fluxes $\left( {\overset{⃑}{J}}_{i}^{m}, {\overset{⃑}{J}}_{i}^{d} \right)$. Based on Equation S3.1, the total transport rate of each ion species ($N_{K^{+}}$, $N_{Cl^{-}}$) was calculated by integrating the normal ${\overset{⃑}{J}}_{i}$, ${\overset{⃑}{J}}_{i}^{m}$ and ${\overset{⃑}{J}}_{i}^{d}$ along each designed boundary around the nanopipette tip (Figure S3.1)^8^.

$$N_{i}=\int_{C} {\overset{⃑}{J}}_{i}ds=\int_{C} {\overset{⃑}{J}}_{i}^{m}ds+\int_{C} {\overset{⃑}{J}}_{i}^{d}ds=N_{i}^{m}+N_{i}^{d}$$

(S3.1),

where *C*: curve (boundary of integration) and *ds*: elementary arc length of the curve *C*.

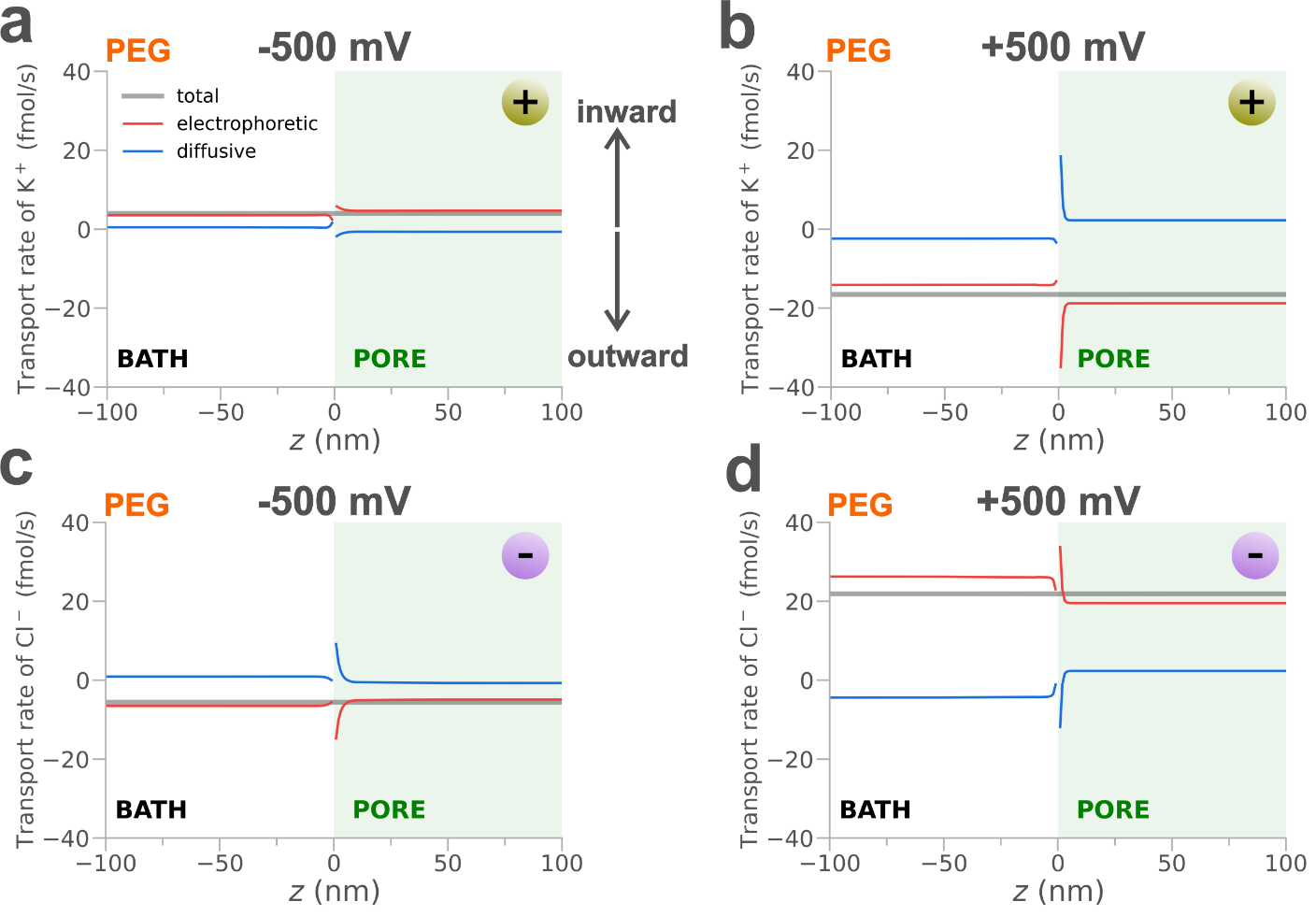

**Figure S3.2** Total, electrophoretic and diffusive transport rates of a) K^+^ for -500 mV, b) K^+^ for 500 mV, Cl^-^ for -500 mV and d) Cl^-^ for -500 mV when PEG is present to the outside bath. The horizontal axis (z) represents the radius in nm of the designed boundaries, as explained in section S3.1. The green positive and purple negative spheres represent K^+^ and Cl^-^, respectively. The inward/outward arrows represent movement of each ion species towards inside/outside the nanopipette tip opening.

*S3.3 Nanopipette sensing region*

Figure S3.3a illustrates the electric potential distribution around the nanopipette tip region when PEG is added in the outer bath. By drawing equipotential lines (each curve represents a constant value for the electric potential in that domain), we noticed that 50% of the applied voltage in the model (ΔV_sens_ = 250 mV) drops within approximately ±20 nm (from the tip interface. This region (40 nm along the z-axis) was defined as the “sensing region” of this system.

In addition, Figure S3.3b shows the distribution of the electric potential $\left( V \right)$ along the z-axis of the geometry for the presence (orange curve) and absence (gray curve) of PEG in the outside solution. In addition, an inset from z = -200 nm to z = 200 nm was included to depict the difference of the “sensing region” in size between the case of PEG and no PEG. To define the “sensing region” for the latter case, we used the equipotential line at -20 nm as the starting point and found the equipotential line inside the pore (120 nm) that reached the same ΔV as in PEG.

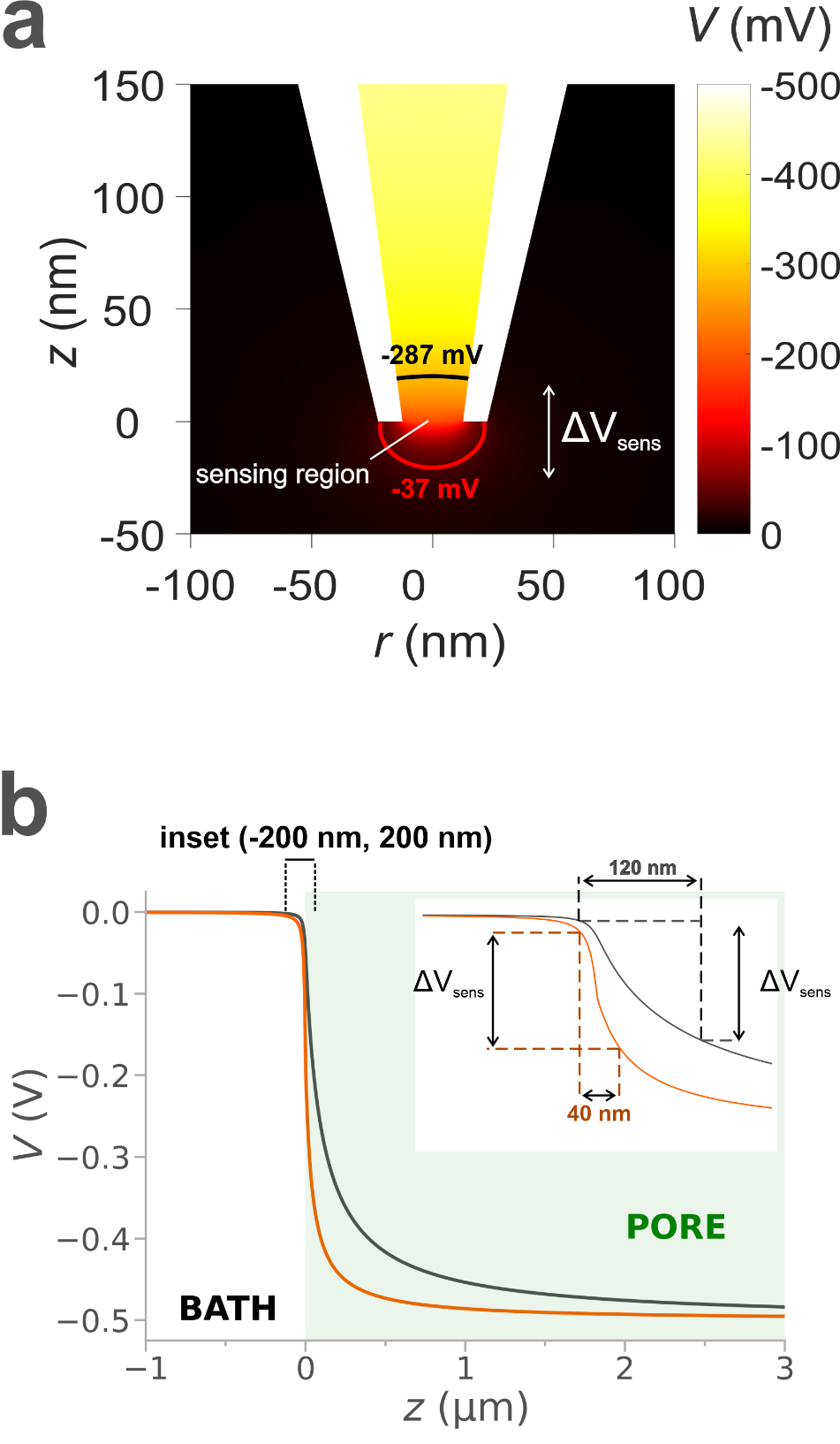

**Figure S3.3** a) Surface colour plot of the electric potential distribution around the nanopipette tip region. The two highlighted curves represent equipotential lines for -287 mV and -37 mV illustrating the “sensing region” where half of the applied voltage drops (ΔV_sens_ = 250 mV) in the presence of PEG. b) Electric potential along the symmetry z-axis of the model geometry (r = 0 nm) for both cases where PEG is present (orange) and absent (grey) from the outside bath. The initial applied voltage is -500 mV at the top boundary of the nanopipette. The inset focuses on the voltage between -200 nm ≤ z ≤ 200 nm to depict the differences between the two “sensing regions” (PEG and no PEG).

*S3.4 Ion transport rates*

Figure S3.4 presents the COMSOL geometry of the finite element model with two additional boundaries that mimic the shape of the equipotential lines for -287 mV and -37 mV at z = ±20 nm, respectively (Figure S3.3a). The surface area enclosed between these two boundaries represents the “sensing region” when PEG is added in the external solution.

To define the transport rates of each ion species in the “sensing region” of the PEG case, we integrated the normal total, electrophoretic and diffusive ion fluxes for K^+^ and Cl^-^ as explained in section S3.2. The integration was performed along the two boundaries mentioned above (z = ±20 nm) and two additional semi-elliptical boundaries (Figure S3.1b) at z = ±5 nm.

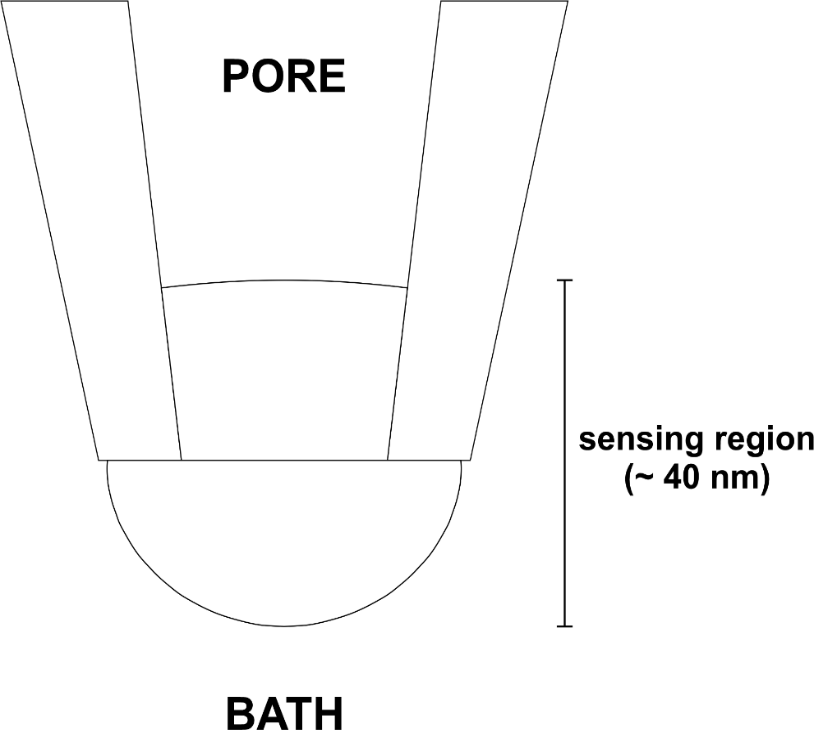

**Figure S3.4** COMSOL geometry design of the finite element model used to calculate the transport rates of each ion species, with the addition of two boundaries identical to the equipotential lines that represented the “sensing region” when PEG is added in the external solution.

The following tables (S3.1, S3.2) present the electrophoretic, diffusive and total transport rate values of potassium cations and chloride anions for ±500 mV with PEG in the external solution. The values at z = ±5 nm, were obtained from the initially designed semi-elliptical boundaries (section S3.1).

**Table S3.1** Electrophoretic $\left( N_{i}^{m} \right)$, diffusive $\left( N_{i}^{d} \right)$ and total $\left( N_{i} \right)$ transport rate of K^+^ and Cl^-^ in [fmol/s] for -500 mV with PEG in the outer solution at the height (z) of 4 equipotential boundaries close to the nanopipette tip. Negative/positive values represent directionality outwards/inwards the nanopipette tip opening.

| **-500 mV** | | | | | | |
| --- | --- | --- | --- | --- | --- | --- |
| **z [nm]** | $\boldsymbol{N}_{\boldsymbol{K}^{\boldsymbol{+}}}^{\boldsymbol{m}}$ **[fmol/s]** | $\boldsymbol{N}_{\boldsymbol{K}^{\boldsymbol{+}}}^{\boldsymbol{d}}$ **[fmol/s]** | $\boldsymbol{N}_{\boldsymbol{K}^{\boldsymbol{+}}}$**[fmol/s]** | $\boldsymbol{N}_{\boldsymbol{Cl}^{\boldsymbol{-}}}^{\boldsymbol{m}}$ **[fmol/s]** | $\boldsymbol{N}_{\boldsymbol{Cl}^{\boldsymbol{-}}}^{\boldsymbol{m}}$ **[fmol/s]** | $\boldsymbol{N}_{\boldsymbol{Cl}^{\boldsymbol{-}}}$ **[fmol/s]** |
| **-20** | 3.53 | 0.49 | 4.02 | -6.53 | 0.93 | -5.60 |
| **-5** | 3.56 | 0.46 | 4.02 | -6.28 | 0.68 | -5.60 |
| **5** | 4.79 | -0.77 | 4.02 | -5.60 | -0.001 | -5.60 |
| **20** | 4.68 | -0.66 | 4.02 | -4.94 | -0.66 | -5.60 |

**Table S3.2** Electrophoretic $\left( N_{i}^{m} \right)$, diffusive $\left( N_{i}^{d} \right)$ and total $\left( N_{i} \right)$ transport rate of K^+^ and Cl^-^ in [fmol/s] for +500 mV with PEG in the outer solution at the height (z) of 4 equipotential boundaries close to the nanopipette tip. Negative/positive values represent directionality outwards/inwards the nanopipette tip opening.

| **+ 500 mV** | | | | | | |
| --- | --- | --- | --- | --- | --- | --- |
| **z [nm]** | $\boldsymbol{N}_{\boldsymbol{K}^{\boldsymbol{+}}}^{\boldsymbol{m}}$ **[fmol/s]** | $\boldsymbol{N}_{\boldsymbol{K}^{\boldsymbol{+}}}^{\boldsymbol{d}}$ **[fmol/s]** | $\boldsymbol{N}_{\boldsymbol{K}^{\boldsymbol{+}}}$**[fmol/s]** | $\boldsymbol{N}_{\boldsymbol{Cl}^{\boldsymbol{-}}}^{\boldsymbol{m}}$ **[fmol/s]** | $\boldsymbol{N}_{\boldsymbol{Cl}^{\boldsymbol{-}}}^{\boldsymbol{m}}$ **[fmol/s]** | $\boldsymbol{N}_{\boldsymbol{Cl}^{\boldsymbol{-}}}$ **[fmol/s]** |
| **-20** | -14.07 | -2.35 | -16.42 | 26.01 | -4.27 | 21.74 |
| **-5** | -14.03 | -2.39 | -16.42 | 25.53 | -3.79 | 21.74 |
| **5** | -18.65 | 2.23 | -16.42 | 19.42 | 2.32 | 21.74 |
| **20** | -18.65 | 2.23 | -16.42 | 19.42 | 2.32 | 21.74 |

**S4	Mechanism of current enhancement upon dsDNA translocation**

*S4.1 Estimating the number of ions carried by single dsDNA in the “sensing region”*

So far, we explained the differences in the baseline current magnitude between the PEG and no PEG models by analysing the ion concentration distributions and transport rates close to the nanopipette tip, especially in the “sensing region”. In this section, we attempt to unravel the mechanism behind the enhanced current magnitude when dsDNA molecules translocate through the nanopipette aperture towards the PEG-enriched bath.

At first, we calculated the number of ions inside the sensing region (Figure S3.4) for both cases (PEG, no PEG). Based on Figure S2.1, the average ion concentration in the sensing region is 52 mM and 102 mM for the PEG and no PEG case, respectively. By surface integration of these values, we obtained the total number of ions for both the PEG and no PEG models as follows:

$$n_{t}={2\pi N}_{A}*\iint r*c_{avg}drdz={\pi N}_{A}*\iint r*\left( c_{K^{+}}+c_{Cl^{-}} \right)drdz$$

(S4.1),

where $N_{A}$: Avogadro’s constant and $c_{K^{+}}, c_{Cl^{-}}$: K^+^ and Cl^-^ concentration, respectively.

We estimate the presence of 972 ions within the sensing reason when PEG is added to the bath compared to 1895 ions when no PEG is added. These values correspond to an ion current at V = -500 mV of -0.92 nA and -3.27 nA, respectively. When single dsDNA molecules translocate through the sensing region towards the outer bath, we observe current peaks of -1.38 nA with PEG and -3.40 nA without PEG. To achieve such values, the number of ions in the sensing region would have to increase by approximately 33% and 4%, respectively. However, we would expect the number of counterions carried by dsDNA molecules in the sensing region to be the same for both case. This is an indication of more complex phenomena associated with the translocation of the molecule and the interface between 0.1 M KCl and 0.1 M KCl with PEG.

*S4.2* *Model for interface displacement due to dsDNA translocation*

We addressed the interaction between single dsDNA molecules translocating through the nanopipette aperture as a displacement of the internal and external solutions interface towards the outside bath. We achieved this by designing a rectangle with a fixed width of 12.5 nm (equal to pore tip radius) and a height ranging from 2 nm to 30 nm, for z < 0 nm (Figure S4.1a, b). The boundary conditions allocated to this new domain were identical to those applied in the nanopipette pore domain. Then, we solved the simulation to find which height provided the closest current value to the experimental translocation peak current (Figure 1c, main text).

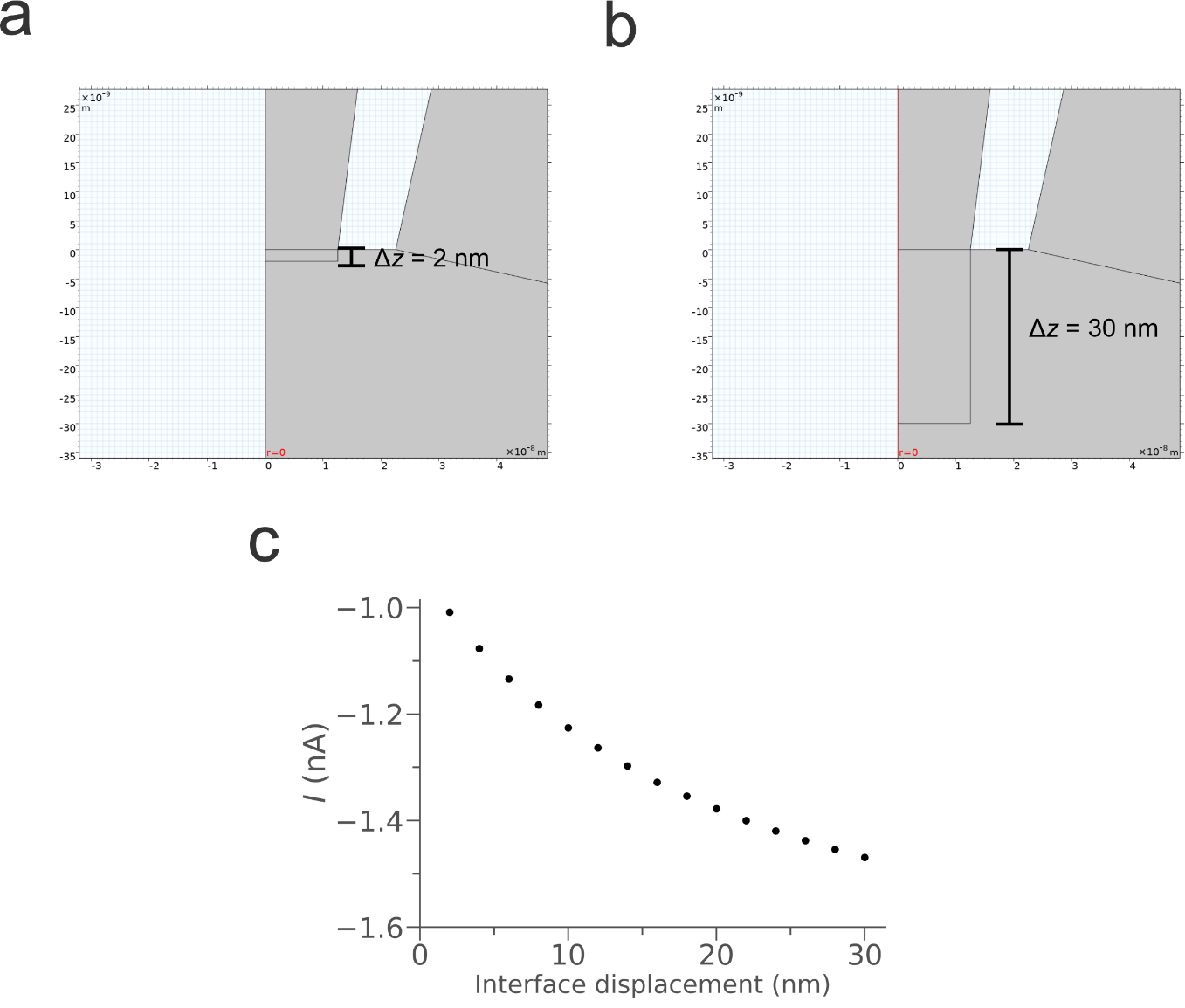

**Figure S4.1** Model for interface displacement towards the bath solution. a, b) Exported COMSOL geometries for the minimum and maximum interface displacement along the z-axis with a fixed width equal to the radius of the nanopipette tip aperture (12.5 nm). c) Simulated current for different interface displacements towards the bath solution.

Depending on the length of the interface displacement, the current peak magnitude increases significantly until 20 nm where it starts to saturate (Figure S4.1d). At z = -16 nm, the simulated current has a magnitude of -1.33 nA which agrees with the experimental value for a 4.8 kbp dsDNA translocating through the nanopipette (Table S4.1).

*S4.3 Effect of dsDNA size on translocation current*

In this section, we investigate the effect of the dsDNA size on the experimentally recorded translocation current. By using the same experimental configuration as shown in Figure 1 in the main text, we recorded a trace of 30 s for every dsDNA molecule size (0.7, 1.5, 2, 3, 4, 4.8 and 7 kbp) where a different nanopipette was used for each size for both PEG and no PEG cases. Figure S4.2a shows a representative event for each current trace with bigger dsDNA molecules corresponding to higher current peak magnitudes. All events per trace are illustrated in the scatter plot in Figure S4.2b in terms of current peak maxima and dwell time, where the latter is the duration of the event calculated at full-width half-maximum. The dsDNA size can be easier discriminated using the current peak maxima values compared to their dwell times.

By calculating the mean current peak maxima and standard deviation for each set of events we observed a dependency between the peak maxima and dsDNA molecule size, as depicted in Figure S4.3c (orange data points). The same trend is not noticeable for the case without PEG in the outer bath (grey data points) with a detection limit of 4.8 kbp.

**Table S4.1** Experimental and simulated translocation current magnitudes for each dsDNA size and interface displacement.

| dsDNA size (kbp) | 0.7 | 1.5 | 2 | 3 | 4 | 4.8 | 7 |
| --- | --- | --- | --- | --- | --- | --- | --- |
| Experimental Current Peak Maxima (nA) | 0.13 | 0.18 | 0.24 | 0.30 | 0.37 | 0.41 | 0.44 |
| Displacement (nm) | **4** | **6** | **8** | **10** | **14** | **16** | **18** |
| Simulated Current Peak Maxima (nA) | 0.16 | 0.21 | 0.26 | 0.30 | 0.38 | 0.41 | 0.43 |

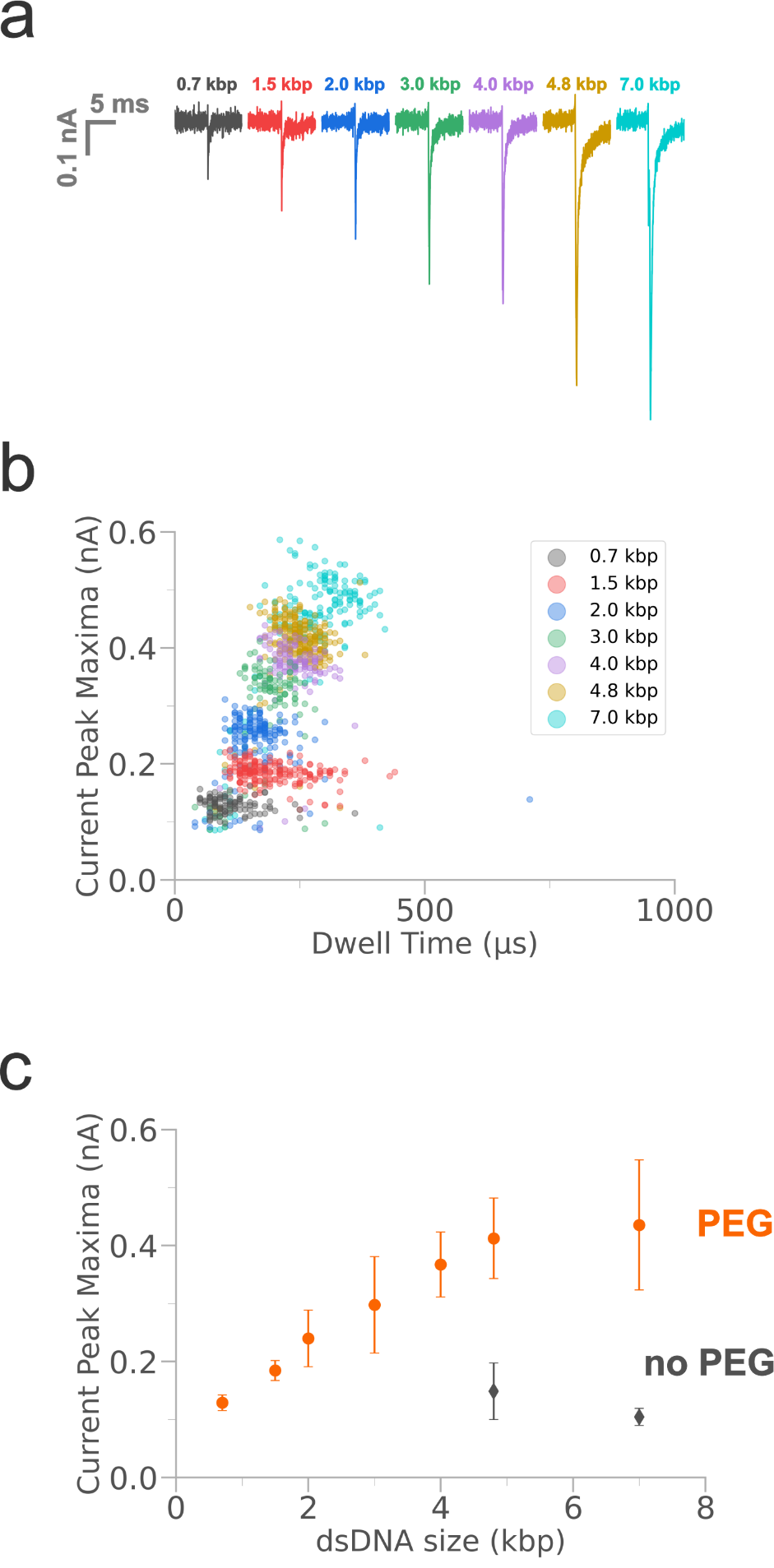

**Figure S4.2** Experimental translocation currents for different dsDNA molecule sizes. a) Representative event for each dsDNA size showing current peaks. b) Scatter plot for all events for each dsDNA size in terms of current peak maxima versus dwell time. c) Mean current peak maxima over size of dsDNA molecules translocating through the nanopipette tip aperture towards the bath with (orange) and without (grey) PEG. The error bars represent the standard deviation.

*S4.4 Effect of an external applied pressure to the nanopipette*

I-V curves were recorded while applying a positive pressure of 1 bar at the back of the nanopipette, which is filled with 0.1 M KCl (without dsDNA) and immersed in 0.1 M KCl with 50% (w/v) PEG 35K^9,10^. As Figure S4.3a illustrates, we observe a different voltametric response when a positive pressure is applied at the back of the nanopipette. The unique *i*-*V* curve associated with the presence of PEG in the external solution with no pressure applied (black curve) is replaced by an *i*-*V* similar to the one reported when PEG was absent from the outer bath (purple curve). This finding indicates that once an external pressure is introduced $\left( p_{app}=1 bar \right)$, convective flow of the internal solution pushes PEG molecules away from the nanopipette tip aperture further in the outer bath. In contrast, under standard pressure conditions $\left( p_{app}=0 bar \right)$ PEG molecules are located in close proximity to the nanopipette tip opening, where they interact with ions in the solution influencing the recorded background current. As a result, when single dsDNA molecules pass through the tip aperture, they temporarily displace PEG molecules to complete their translocation to the outside bath which causes an interface displacement and ion reorganization leading to the enhanced translocation current peak measure in the presence of PEG. Figure S4.3b shows the experimental current trace recorded during dsDNA (4.8 kbp) translocation before, during and after the application of pressure $\left( p_{app}=1 bar \right)$. The translocation current enhancement due to the presence of PEG is completely nullified when the pressure is applied and it comes back to normal when the pressure is released.

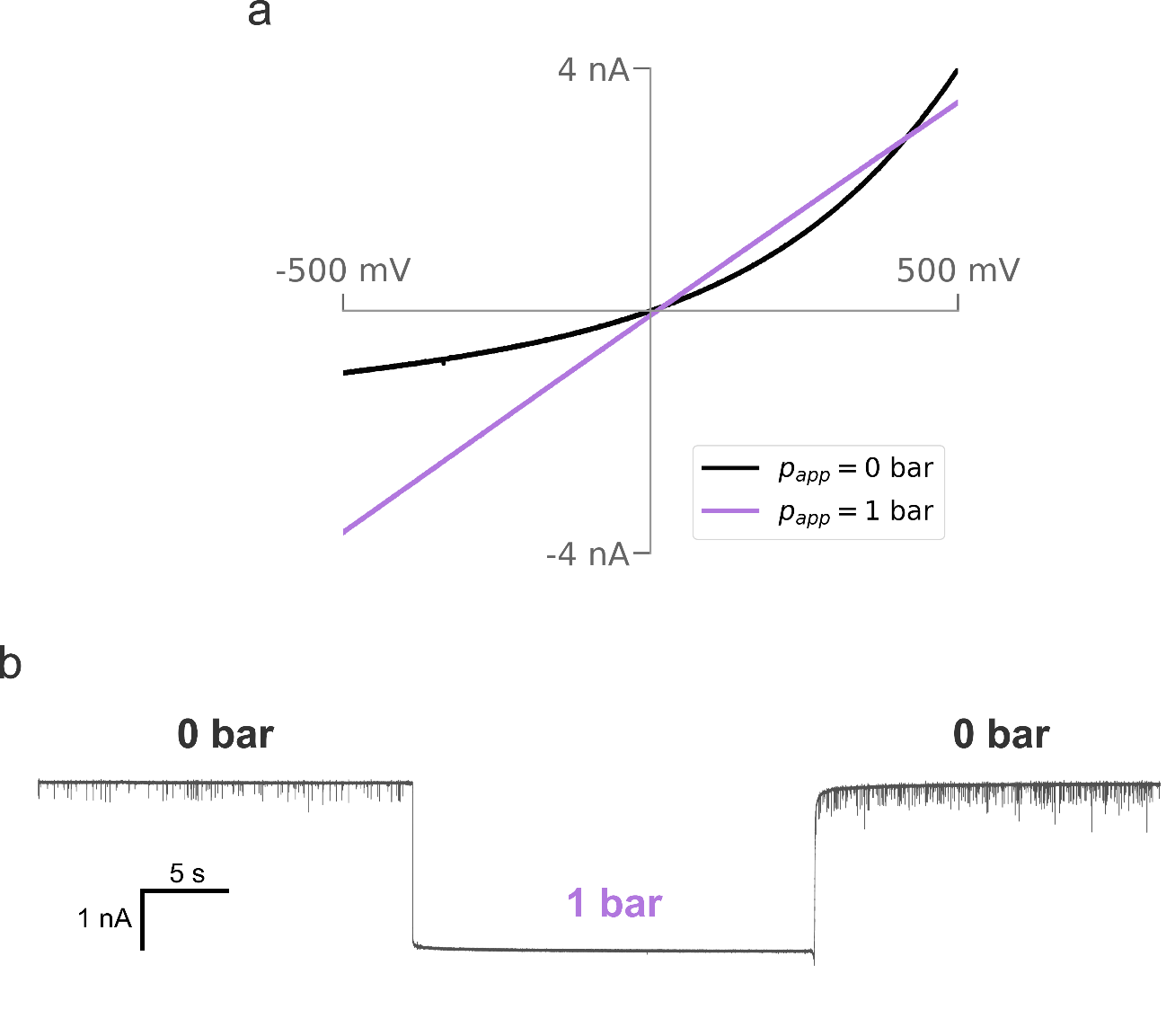

**Figure S4.3** a) Experimental voltammogram for V = ±500 mV, when a pressure of 0 and 1 bar (black and purple curves, respectively) is applied at the back of the nanopipette filled with 0.1 M KCl and immersed in 0.1 M KCl with 50% (w/v) PEG 35K. b) Experimental current trace during 4.8 kbp dsDNA translocation before, during and after the application of 1 bar pressure at the back of the nanopipette for V = -500 mV.
